# m^6^A-Driven Intratumoral Cholesterol Biosynthesis Fuels Castration-Resistant Prostate Cancer Progression

**DOI:** 10.64898/2026.09.24.753979

**Authors:** Ji Hoon Lee, Zhao Zhang, Juyeong Hong, Ji-Hoon Jeong, Huihui Ye, Divyaa Srinivasan, Lei Kang, Santosh Lamichhane, Nicolas Amselle, Kayode A. John, Matthew T. Hinds, Apeksha N. Agarwal, Dong Min Lee, Songmi Han, Martine P. Roudier, Hung-Ming Lam, Eva Corey, Kexin Xu

## Abstract

Both nuclear pore complexes (NPCs) and RNA *N*^6^-methyladenosine (m^6^A) machinery are indispensable for proper cellular function. Although their collaborative roles in the nuclear export of messenger RNAs (mRNAs) have been reported, it remains ambiguous whether and how this collaboration may contribute to cancer progression. Here we identify a functional cooperation between NPCs and m^6^A signaling that promotes the development of castration-resistant prostate cancer (CRPC). We showed that nuclear export of m^6^A-modified mRNAs, mediated by the interaction between RNA methyltransferase METTL3 and the nucleoporin NUP93, is functionally coupled to cholesterol biosynthesis. Given that cholesterol-fueled intratumoral androgen production is one of the mechanisms driving CRPC, we demonstrated that overexpression of the wild-type METTL3 or NUP93, but neither the enzymatically dead METTL3 nor the mutant NUP93 that loses METTL3-interacting capability, elevates intracellular levels of androgens, activates AR signaling under castrate condition, and promotes androgen-independent growth of prostate cancer cells both *in vitro* and *in vivo*. Importantly, pharmacological inhibition of METTL3 or targeted demethylation on mRNAs encoding key cholesterol biosynthesis enzymes effectively suppressed CRPC malignancy. Together, these findings uncover a therapeutically targetable m^6^A-METTL3-NUP93 axis that links nuclear mRNA export and metabolic reprogramming to fuel CRPC progression, providing a conceptually new strategy for the treatment of this lethal disease.

## Introduction

As the gatekeeper of nuclear-cytoplasmic transport, nuclear pore complexes (NPCs) play critical roles in regulating a plethora of biological processes that are essential for cells, and when malfunctioning, are associated with many human diseases (1,2). For example, numbers of NPCs are often found to be increased in cancer cells, suggesting an increased reliance on nuclear transport machinery (3,4). Function and assembly of NPCs are executed by around 30 different proteins called nucleoporins (NUPs) (5). Based on sequence conservation and functional distinction, NUPs are classified into three classes: “structural NUPs” that build NPC architecture; “membrane NUPs” that anchor the whole complex to the nuclear envelope; and “FG-NUPs” that control the nucleocytoplasmic trafficking of macromolecules like messenger RNA (mRNAs) and proteins (6). Several NUPs have been linked to cancers through chromosomal translocations (7,8), single point mutations (9) and altered expression levels (10,11). However, considering that NPCs transport a diversity of cargoes and that NUPs have functions independent of transport process (12,13), it is still unsettled whether and how dysregulation of NUPs disrupts normal functioning of NPCs and subsequently leads to cancer progression.

Several studies demonstrate that RNA methylation at the *N*^6^-position of adenosine (m^6^A) regulates the nuclear export of selective mRNA molecules across NPCs (14–16). Methyl groups are deposited to adenosine within the DRACH consensus motif (D=A/G/U, R=A/G, H=A/C/U) of mRNAs by a multiprotein methyltransferase complex consisting of the catalytic subunit METTL3 and its partners like METTL14 and WTAP (17,18). The modification can also be removed by RNA demethylases, FTO and ALKBH5 (19,20). Accumulating evidence suggests that dysregulation of m^6^A modification leads to aggressive phenotypes in cancer cells, such as metastasis, therapeutic resistance, immunosuppression, etc (21). However, the underlying mechanisms are highly diverse and cancer-type specific. It has been reported that some of m^6^A regulators interact with export receptors as well as adaptor proteins that mediate the transfer of mRNA cargos through NPCs to the cytoplasm (15,16,22). Previously, we uncovered that METTL3 can be localized on NPCs through direct association with the nucleoporin protein NUP93 (23). The METTL3-NUP93 interaction facilitates the nuclear export of m^6^A-modified mRNAs through nuclear pores. Despite the well demonstrated role of m^6^A modification in cooperating with NPCs to export nuclear mRNAs, no studies have ever elucidated whether and how the crosstalk between m^6^A machinery and NPCs contributes to cancer progression.

Here we uncovered an important contribution of the NPC-m^6^A interplay to the development of castration-resistant prostate cancer (CRPC), a lethal form of the disease (24). Via integrative analysis of global m^6^A landscape and gene expression patterns in the nuclear compartment of prostate cancer cells, we identified a group of m^6^A-modified mRNAs whose nuclear export requires an intact METTL3-NUP93 complex. Intriguingly, these mRNAs are functionally enriched in biosynthesis of cholesterol, which plays an essential role in steroidogenesis within prostate tumors and accelerates disease progression under castrate conditions (25–30). Indeed, prostate cancer cells with the operative m^6^A-METTL3-NUP93 axis acquired castration-resistant phenotypes. Most importantly, we demonstrated that both pharmacological disruption of m^6^A signaling and targeting methylated sites on mRNAs for cholesterol synthesis using RNA-editing tools can lead to inhibition of CRPC, highlighting the discoveries of novel therapeutic strategies for advanced prostate cancer.

### NUP93 is highly upregulated and functionally required in castration-resistant prostate cancer

A growing body of evidence shows that several nucleoporins, the key components of nuclear pore complex (NPC), are dysregulated across human cancers and have emerged as critical regulators of tumor cell fitness (31–35). To determine whether NPC dysfunction contributes to prostate cancer progression, we systematically examined the expression and essentiality of all nucleoporin genes that are available in TCGA and DepMap for prostate cancer, respectively. One of the nucleoporin genes called NUP93 was selected for further characterization for the following reasons. First, NUP93 is one of the top nucleoporin genes that are significantly upregulated in prostate cancer compared to normal tissues (**Fig. 1A**). We further confirmed higher expression of NUP93 in prostate cancer than in the matched normal counterparts in TCGA dataset (**Supplemental Fig. 1A**) and several independent prostate cancer cohorts (**Supplemental Fig. 1B**). Second, NUP93 is the most essential nucleoporin gene in two prostate cancer cell lines (**Fig. 1B**), and indeed when we knocked down the top four nucleoporin genes that are upregulated in prostate cancer, NUP93 depletion showed the most significant growth inhibition in both LNCaP and C4-2B cells (**Fig. 1C**). Finally, our recent work has uncovered an impactful interaction between NUP93 and METTL3 (23), which facilitates the export of m^6^A-modified mRNAs through the nuclear pore complexes. To demonstrate the clinical relevance of this interaction in prostate cancer, we performed immunohistochemical analysis of both NUP93 and METTL3 in tissue microarrays (TMA) containing prostate tumors with matched benign tissues and CRPC samples. 51.1% of CRPC cases express high levels of both proteins in contrast to 45.6% in primary prostate tumors and 33.6% in normal adjacent tissues (**Fig. 1D**). Concurrent high expression of both proteins was the predominant expression pattern in CRPC specimens, occurring noticeably more frequently than high expression of either protein alone or low expression of both (**Fig. 1E**). To further verify its functional importance in prostate cancer cells, we depleted NUP93 in androgen-dependent (LNCaP, DuCaP) and castration-resistant (C4-2B, 22Rv1) prostate cancer cell lines. NUP93 depletion consistently impaired proliferation across all models (**Fig. 1F-H** and **Supplemental Fig. 1C-D**) and significantly reduced migration and invasion in CRPC cells (**Fig. 1I-J**).

**Figure 1.**
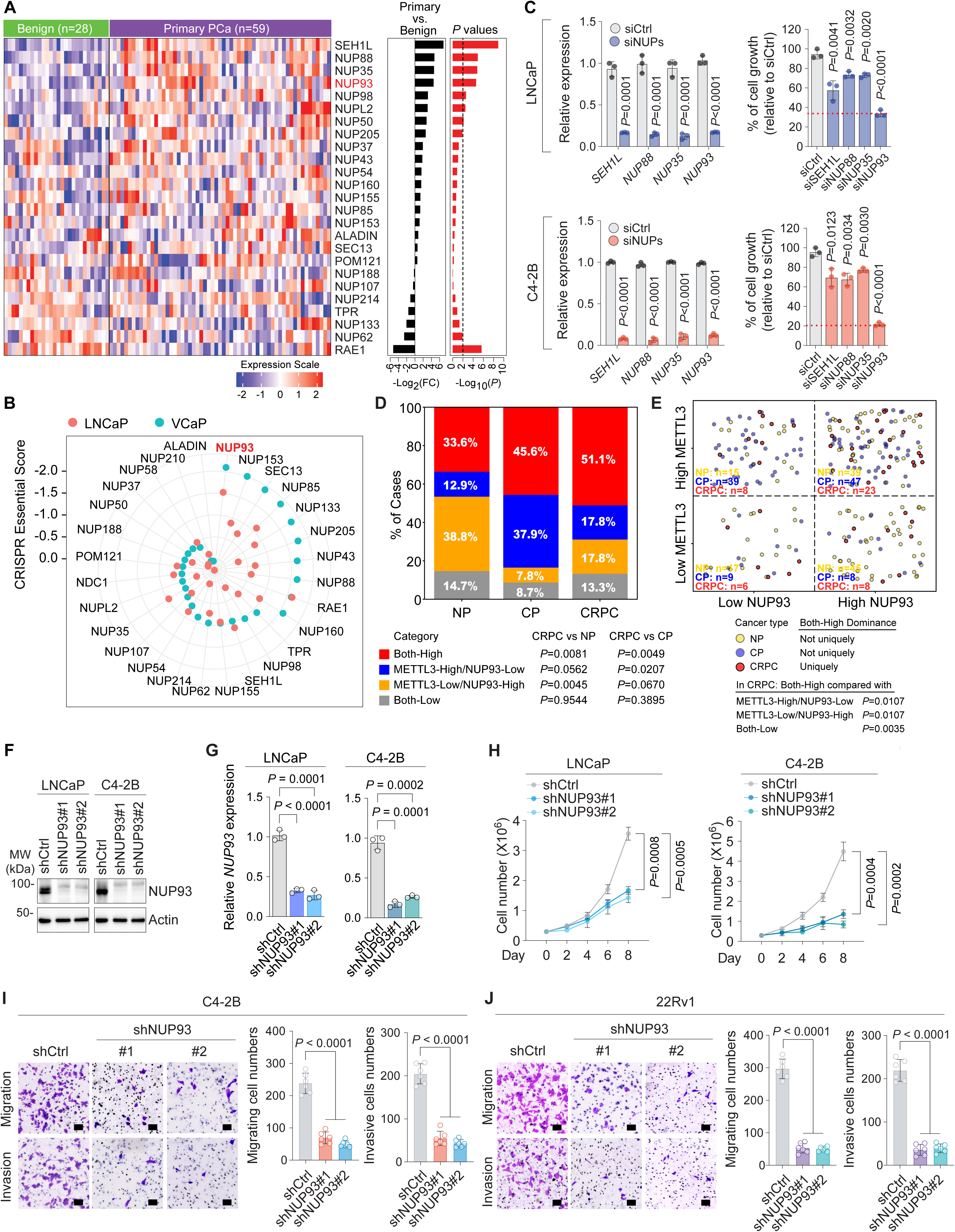
NUP93 is upregulated and required in advanced prostate cancer. **(A)** Pan-cancer or prostate-focused analysis of nucleoporin gene expression comparing prostate tumors with nonmalignant prostate tissues in GSE35988. **(B)** DepMap dependency scores for nucleoporin genes in LNCaP and C4-2B cells; NUP93 is highlighted. **(C)** RT-qPCR analysis of the indicated nucleoporin genes (left) and cell growth analysis (right) in LNCaP and C4-2B cells following depletion of the four nucleoporins most highly upregulated in prostate cancer. Gene expression and cell growth were normalized to those in cells transfected with control siRNA (siCtrl). **(D)** Percentages of the cases in tumor-adjacent nonmalignant prostate (NP), primary prostate cancer (CP), and castration-resistant prostate cancer (CRPC) samples according to the combination of METTL3 and NUP93 protein levels. Specimens were classified as high levels of both proteins (Both-High), high level of one but low of the other (METTL3-High/NUP93-Low and METTL3-Low/NUP93-High), and low levels of both proteins (Both-Low). Bars show the percentage of specimens in each protein expression category. *P* values were determined by two-sided Fisher’s exact tests comparing the proportion of cases in each protein expression category between CRPC and NP or CP. **(E)** Distribution of individual specimen according to the combination of METTL3 and NUP93 protein levels as defined in **(D)**. NUP93 expression status is shown on the X-axis and METTL3 on the Y-axis. Each circle represents one tissue sample, colored by the disease group and randomly positioned with the corresponding quadrant for visualization. Sample numbers of tumor-adjacent nonmalignant prostate (NP), primary prostate cancer (CP), and castration-resistant prostate cancer (CRPC) in each quadrant were listed. *P* values were determined by two-sided Fisher’s exact tests comparing the frequency of each protein expression category in CRPC with NP or CP. **(F, G)** Validation of NUP93 depletion in LNCaP and C4-2B cells expressing control (shCtrl) or NUP93-targeting shRNAs (shNUP93#1 and shNUP93#2). NUP93 protein and mRNA levels were assessed by immunoblotting **(F)** and RT-qPCR **(G)**, respectively. Actin was served as the loading control for immunoblotting. *NUP93* mRNA expression was normalized to *GAPDH* and is presented relative to the corresponding control cells. **(H)** Proliferation of LNCaP and C4-2B cells expressing control (shCtrl) or NUP93-targeting shRNAs (shNUP93#1 and shNUP93#2). **(I, J)** *In vitro* transwell migration and invasion assays in C4-2B **(I)** and 22Rv1 **(J)** cells expressing control (shCtrl) or NUP93-targeting shRNAs (shNUP93#1 and shNUP93#2). Representative images of migration (top) and invasion (bottom) are shown on the left, with the corresponding quantification shown on the right. Scale bars, 100 μm. Data are mean ± SD. n = 3 independent experiments or samples. *P* values in **(G-J)** were determined by two-tailed paired *t* tests.

In summary, we identify NUP93 as a clinically relevant and functionally essential NPC component in advanced prostate cancer, and our work has provided a rationale to investigate the molecular mechanisms by which NUP93 promotes disease progression via interaction with METTL3.

### NUP93 cooperates with METTL3 and requires METTL3 binding to support prostate cancer progression

Our previous work discovered a functional complex between METTL3 and NUP93, which plays a critical role in m^6^A-dependent nuclear export of select mRNA species (23). Here, we speculated that the intact METTL3-NUP93 complex is required for prostate cancer malignancy. We first examined whether the METTL3-NUP93 interaction is preserved in prostate cancer. Co-immunoprecipitation (Co-IP) revealed a robust interaction between endogenous METTL3 and NUP93 in both androgen-dependent (LNCaP, DuCaP) and castration-resistant (C4-2B, 22Rv1) prostate cancer cells (**Fig. 2A**), demonstrating the common existence of the METTL3-NUP93 complex in prostate cancer cells. We next assessed the importance of interaction with METTL3 in mediating the oncogenic activity of NUP93 in prostate cancer cells. Using our previously characterized NUP93 mutant (A333P) that is defective in binding with METTL3 (23), we first confirmed by proximity ligation assay (PLA) that the A333P substitution disrupts the METTL3-NUP93 interaction at the nuclear envelope in prostate cancer cells (**Fig. 2B**). We then performed rescue experiments that replace the endogenous NUP93 in prostate cancer cells with either the wild-type NUP93 (hereafter N93-WT) or the A333P mutant that loses interaction with METTL3 (hereafter N93-AP). Knockdown of NUP93 significantly blocked proliferation, anchorage-independent colony formation, migration, and invasion in prostate cancer cells, yet overexpression of NUP93-WT fully restored, whereas NUP93-AP failed to rescue these phenotypes (**Fig. 2C-H** and **Supplemental Fig. 2A**). It is worthy of note that METTL3 protein abundance was not changed in the NUP93 rescue system, suggesting that the tumor-promoting activity of NUP93 is dependent on association with METTL3 rather than regulation of METTL3 protein levels. Finally, we evaluated the effects of NUP93 and its interaction with METTL3 on prostate cancer *in vivo* (**Figure 2I-J**). NUP93 depletion significantly retarded tumor growth in the xenograft mouse model of CRPC, which can be only rescued by re-expression of NUP93-WT but not NUP93-A333P, further confirming that the oncogenic activity of NUP93 in prostate cancer requires functional cooperation with METTL3.

**Figure 2.**
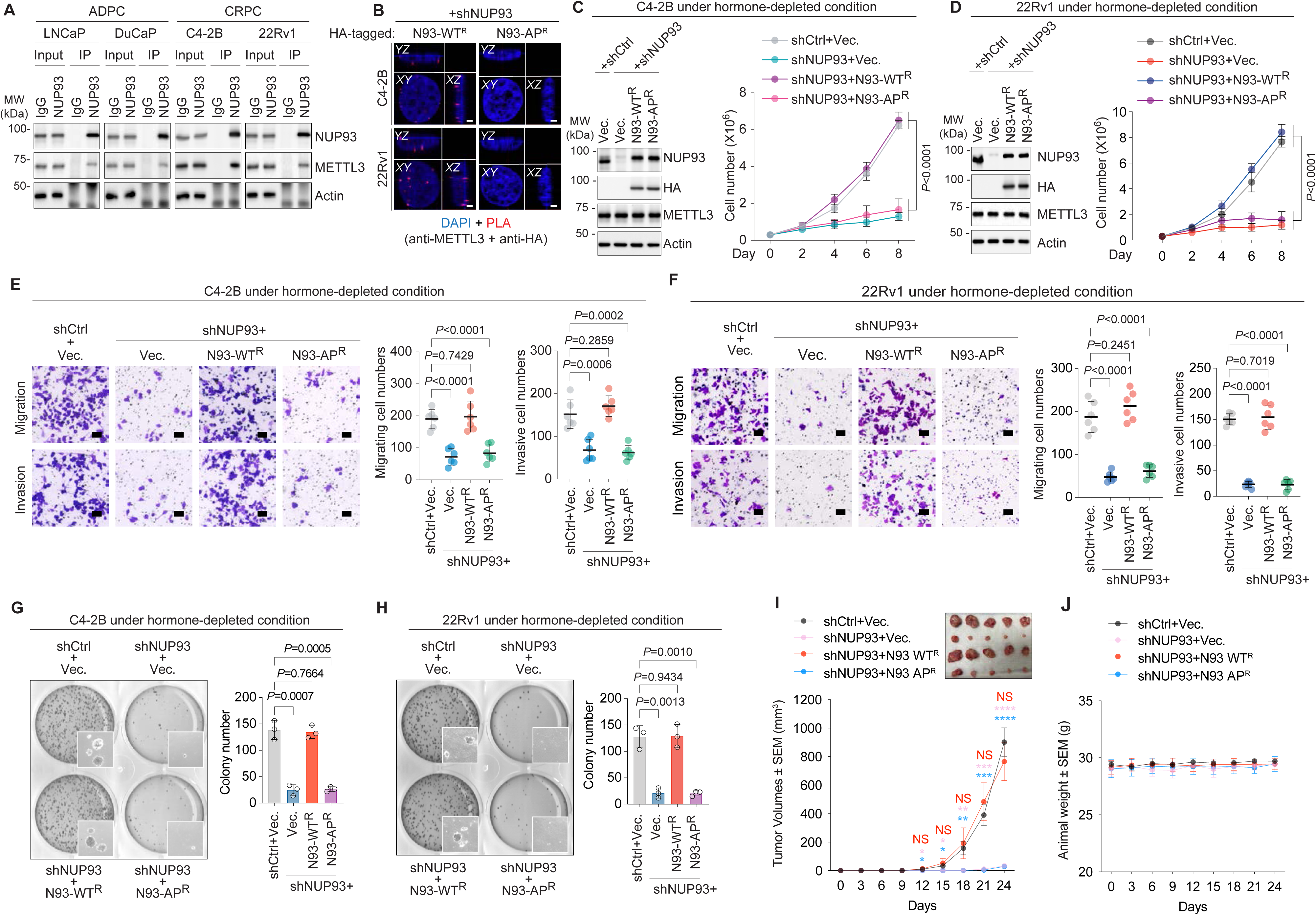
The tumor-promoting activity of NUP93 requires its interaction with METTL3. **(A)** Endogenous co-immunoprecipitation (Co-IP) of METTL3 and NUP93 in androgen-dependent prostate cancer (ADPC; LNCaP and DuCaP) and castration-resistant prostate cancer (CRPC; C4-2B and 22Rv1) cells. Blots are representative of three independent experiments. **(B)** Proximity ligation assay (PLA) detecting METTL3-NUP93 association in C4-2B and 22Rv1 cells expressing NUP93-WT (N93-WT) or the METTL3-binding-defective NUP93-A333P mutant (N93-AP) using anti-METTL3 and anti-HA antibodies. Nuclei are stained with DAPI. Representative images are shown. Scale bars, 2 μm. **(C, D)** Rescue of cell proliferation following endogenous NUP93 depletion and re-expression of HA-tagged wild-type NUP93 (N93-WT) or the METTL3-binding-defective NUP93-A333P mutant (N93-AP) in C4-2B **(C)** and 22Rv1 **(D)** cells. Immunoblot analysis of HA-tagged NUP93, total NUP93, and METTL3 is shown on the left, with Actin serving as the loading control. Cell proliferation is shown on the right. **(E, F)** *In vitro* transwell migration and invasion assays following endogenous NUP93 depletion and re-expression of NUP93-WT or NUP93-AP in C4-2B **(E)** and 22Rv1 **(F)** cells. Representative images of migrating cells (top) and invading cells (bottom) are shown on the left, with the corresponding quantification shown on the right. Scale bars, 100 μm. **(G, H)** Anchorage-independent colony-formation assays following endogenous NUP93 depletion and re-expression of NUP93-WT or NUP93-AP in C4-2B **(G)** and 22Rv1 **(H)** cells. Representative images are shown on the left, with the corresponding quantification shown on the right. **(I, J)** Xenograft growth of 22Rv1 cells expressing control shRNA (shCtrl), NUP93-targeting shRNA (shNUP93), or shNUP93 together with NUP93-WT or NUP93-AP in castrated male nude mice. Representative tumors and tumor growth curves are shown in **(I)**. Changes in mouse body weight during the experiment are shown in **(J)**. Data are presented as mean ± SD in **(C-H)** and mean ± SEM in **(I, J)**. Data in **(C-H)** are from three independent experiments. 5 mice were included in each group in (I, J). *P* values were determined by two-tailed paired *t* tests.

In addition, we asked whether the methyltransferase activity of METTL3 is essential for the malignant phenotypes in prostate cancer. Substitution of the endogenous METTL3 with the wild-type enzyme (hence M3-WT), but not the catalytically dead mutant (hence M3-CD), restored proliferation and invasive capacity of prostate cancer cells following METTL3 knockdown (**Supplemental Fig. 2B-H**). Taken together, these results indicate that NUP93 acts as a critical oncogenic factor in prostate cancer, which is dependent on its association with METTL3 and integrity of m^6^A signaling, highlighting an important role of m^6^A-METTL3-NUP93 axis in prostate cancer progression.

### METTL3-NUP93 cooperation drives m^6^A-dependent nuclear export of cholesterol biosynthesis transcripts in CRPC

Since our prior work has well demonstrated that the m^6^A-METTL3-NUP93 axis facilitates the nuclear mRNA export (23), we asked whether the mRNA cargos whose nuclear transport requires this axis contribute to its role in prostate cancer progression. Therefore, we performed fractionation RNA-seq and integrated the data with transcriptome-wide m^6^A profile in prostate cancer cells. Silencing either METTL3 or NUP93 induced a remarkable nuclear retention of m^6^A-modified transcripts without altering their total RNA abundance (**Fig. 3A-B**), indicating a post-transcriptional defect in RNA export rather than transcriptional repression. Importantly, there is a significant overlap between the groups of mRNAs that are retained in the nuclear compartments upon METTL3 or NUP93 knockdown (**Fig. 3C**), implying that METTL3 and NUP93 coregulate a common set of mRNAs on their nuclear export. Even more exciting, these overlapped mRNAs are functionally enriched in androgen-responsive programs and cholesterol homeostasis pathways (**Fig. 3D**), both of which have been well approved to drive CRPC progression.

**Figure 3.**
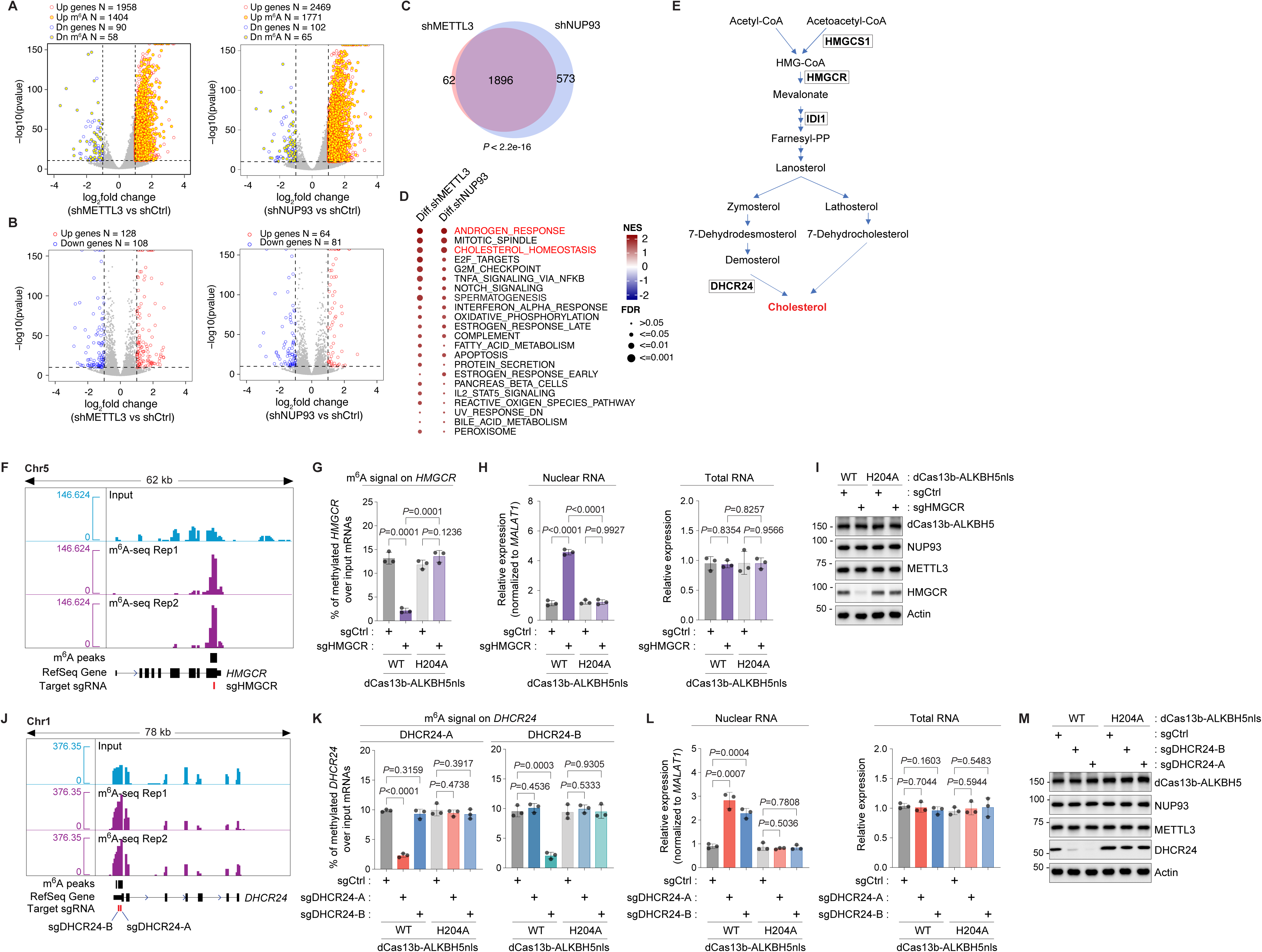
METTL3-NUP93 cooperation promotes m^6^A-dependent nuclear export of cholesterol-biosynthesis transcripts. **(A, B)** Volcano plots showing differentially expressed genes identified by nuclear RNA-seq **(A)** and total RNA-seq **(B)** following METTL3 depletion (left) or NUP93 depletion (right) in LNCaP cells. Red and blue circles indicate significantly upregulated and downregulated genes, respectively, and yellow-filled centers identify m^6^A-modified transcripts. Dashed lines indicate the thresholds of |log_2_fold change| ≥ 1 and adjust *P* values < 1 × 10⁻^10^. **(C)** Venn diagram showing the overlap between genes upregulated in the nuclear RNA fraction following METTL3 or NUP93 depletion. A total of 1,896 genes were commonly upregulated following depletion of either METTL3 or NUP93. *P* < 2.2 × 10⁻^16^, determined by a hypergeometric test. **(D)** Hallmark gene-set enrichment analysis of genes upregulated in the nuclear RNA fraction following METTL3 or NUP93 depletion. Circle color represents the normalized enrichment score (NES), and circle size represents the false discovery rate (FDR). **(E)** Schematic representation of the cholesterol biosynthesis pathway. HMGCS1, HMGCR, IDI1, and DHCR24 are highlighted as cholesterol biosynthesis enzymes encoded by m^6^A-modified transcripts that are accumulated in the nuclear fraction following METTL3 or NUP93 depletion. **(F)** Genome-browser tracks showing input and m^6^A-seq signals from two biological replicates across the *HMGCR* locus. The identified m^6^A peak and the target site of sgHMGCR used for dCas13b-ALKBH5nls (nuclear localization signal)-mediated site-specific RNA demethylation are indicated. **(G)** m^6^A enrichment on *HMGCR* mRNA in cells expressing wild-type dCas13b-ALKBH5nls (WT) or the catalytically inactive H204A mutant together with control sgRNA (sgCtrl) or HMGCR-targeting sgRNA (sgHMGCR). m^6^A enrichment was measured by m^6^A-RIP-qPCR and is presented as the percentage of methylated *HMGCR* RNA relative to input. **(H)** Nuclear (left) and total (right) *HMGCR* mRNA levels in cells expressing wild-type dCas13b-ALKBH5nls or the H204A mutant together with sgCtrl or sgHMGCR. Nuclear *HMGCR* mRNA was normalized to *MALAT1*, and total *HMGCR* mRNA was normalized to *GAPDH*. **(I)** Immunoblotting analysis of dCas13b-ALKBH5nls, NUP93, METTL3, and HMGCR under the conditions described in (G). Actin was served as the loading control. Blots are representative of three independent experiments. **(J)** Genome-browser tracks showing input and m^6^A-seq signals from two biological replicates across the *DHCR24* locus. The identified m^6^A peaks and the distinct target sites of sgDHCR24-A and sgDHCR24-B used for dCas13b-ALKBH5nls-mediated site-specific RNA demethylation are indicated. **(K)** m^6^A enrichment at the DHCR24-A (left) and DHCR24-B (right) target regions in cells expressing wild-type dCas13b-ALKBH5nls (WT) or the catalytically inactive H204A mutant together with sgCtrl, sgDHCR24-A, or sgDHCR24-B. m^6^A enrichment was measured by m^6^A-RIP-qPCR and is presented as the percentage of methylated *DHCR24* RNA relative to input. **(L)** Nuclear (left) and total (right) *DHCR24* mRNA levels in cells expressing wild-type dCas13b-ALKBH5-NLS or the H204A mutant together with sgCtrl, sgDHCR24-A, or sgDHCR24-B. Nuclear *DHCR24* RNA was normalized to *MALAT1*, and total *DHCR24* RNA was normalized to *GAPDH*. **(M)** Immunoblotting analysis of dCas13b-ALKBH5nls, NUP93, METTL3, and DHCR24 under the conditions described in **(K)**. Actin served as the loading control. Blots are representative of three independent experiments. Data in (G-H and K-L) are presented as mean ± SD from three independent experiments. *P* values were determined by two-tailed paired *t* tests for the indicated comparisons.

Cellular fractionation-qPCR analysis further confirmed that NUP93 or METTL3 knockdown selectively impaired the nuclear export of key transcripts encoding cholesterol biosynthesis enzymes (*HMGCR*, *HMGCS1*, *IDI1*, *DHCR24*) (**Fig. 3E**), while leaving their total mRNA levels unchanged. This effect could be fully restored by re-expression of the wild-type NUP93 (**Supplemental Fig. 3A-D**) or METTL3 (**Supplemental Fig. 3E-H**), but neither METTL3-binding-defective NUP93 mutant (N93-AP) nor the catalytically inactive METTL3 mutant (M3-CD). These data suggest that METTL3-mediated m^6^A deposition and the METTL3-NUP93 complex cooperate to ensure efficient nuclear transport of the mRNAs critical for cholesterol biosynthesis and metabolism, one of the key mechanisms driving CRPC.

Furthermore, we performed site-specific RNA demethylation using a dCas13b-ALKBH5nls system to directly test whether m^6^A signal is required for efficient nuclear export of these transcripts (23). Targeted sgRNAs were designed to remove the m^6^A signals on *HMGCR* (**Fig. 3F**) and *DHCR24* (**Fig. 3J**) mRNAs according to our global m^6^A profiling data. Compared to the control sgRNA and enzymatically dead mutant (H204A), co-expression of the targeted sgRNAs and wild-type ALKBH5 significantly decreased m^6^A intensities on both transcripts (**Fig. 3G** and **3K**), which is coincident with nuclear retention of these mRNAs (**Fig. 3H** and **3L**) and reduction in their protein levels (**Fig. 3I** and **3M**). There are no changes in total mRNA expression, and METTL3 or NUP93 protein levels are constant in the dCas13b-ALKBH5nls system. All these data validated that m^6^A modification guarantees nuclear export and proper expression of these mRNAs involved in cholesterol biosynthesis.

### METTL3-NUP93 axis promotes cholesterol accumulation and intratumoral androgen synthesis in CRPC

Since several mRNAs encoding cholesterol biosynthesis enzymes depends on the integrity of m^6^A-METTL3-NUP93 axis for their nuclear export, we asked whether disruption of this axis changes cholesterol levels in prostate cancer cells. Knockdown of METTL3 or NUP93 markedly reduced cellular cholesterol levels across multiple CRPC cell lines when maintained under androgen-deprived and fat-free conditions, and re-expression of the wild-type METTL3 or NUP93 restored cholesterol abundance (**Fig. 4A-B**). In contrast, the catalytically dead METTL3 (M3-CD) and METTL3-binding-deficient NUP93 mutant (N93-AP) failed to rescue cholesterol levels, confirming that m^6^A modification and the METTL3-NUP93 interaction are both required for maintaining cholesterol amounts in CRPC cells. Moreover, we employed the dCas13b-based tool to selectively increase the on-site m^6^A intensities on *HMGCR*, one of the target mRNAs of the m^6^A-METTL3-NUP93 axis, (**Supplemental Fig. 4A**). *HMGCR* encodes the rate-limiting enzyme of the cholesterol-producing mevalonate pathway (28), and we found enhanced nuclear export of *HMGCR* mRNAs only when both target sgRNA and wild-type METTL3 were co-expressed with no change in total mRNA levels (**Supplemental Fig. 4B**), which leads to increased HMGCR protein levels and cholesterol amounts in prostate cancer cells (**Fig. 4C**).

**Figure 4.**
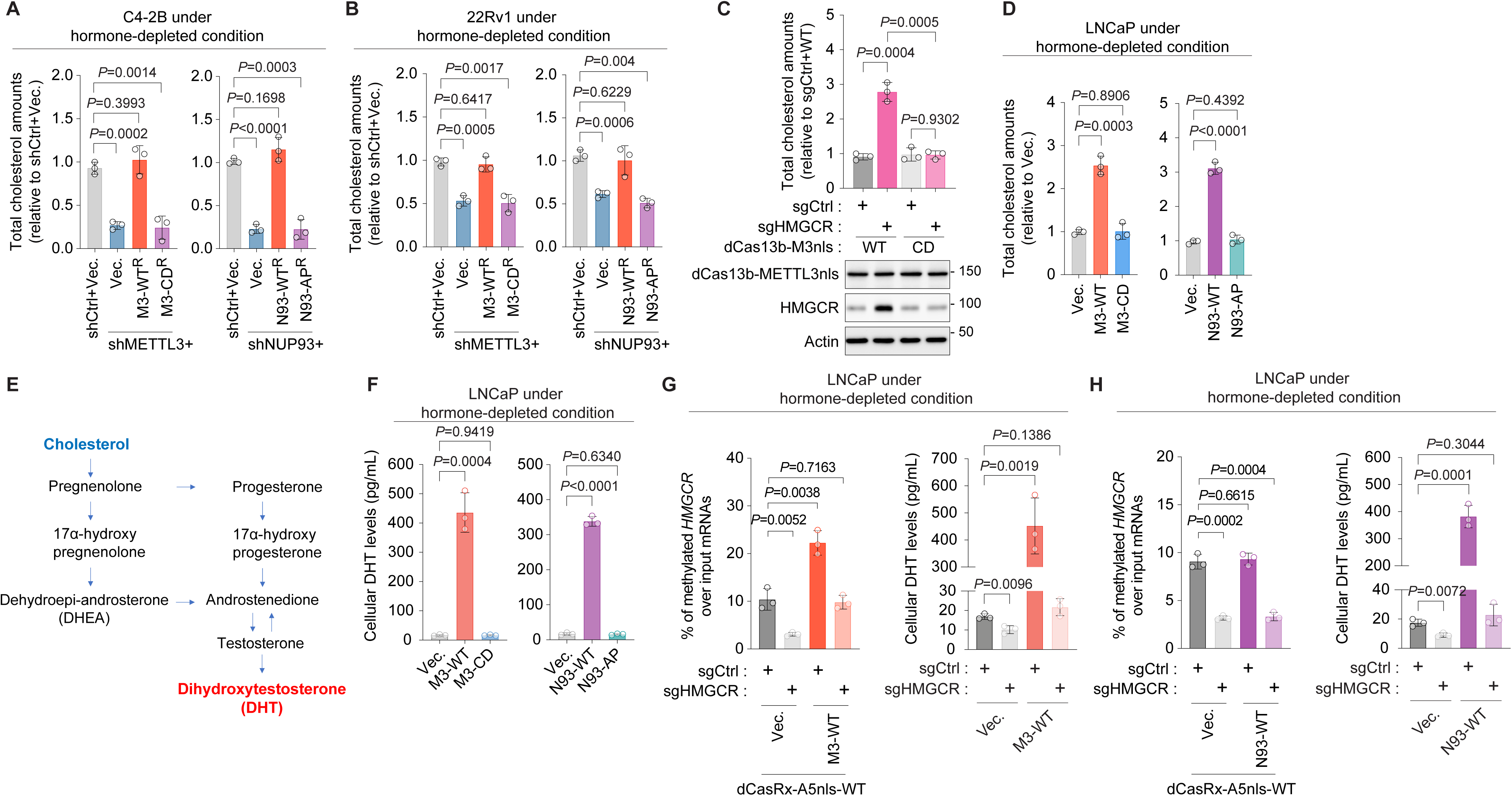
The m^6^A-METTL3-NUP93 axis increases cholesterol abundance and androgen synthesis under hormone-depleted conditions. **(A, B)** Total cellular cholesterol levels in C4-2B (A) and 22Rv1 (B) cells following depletion of endogenous METTL3 or NUP93 and re-expression of wild-type METTL3 (M3-WT), catalytically inactive METTL3 (M3-CD), wild-type NUP93 (N93-WT), or the METTL3-binding-defective NUP93 mutant (N93-AP). Cells were maintained under hormone-depleted conditions. Cholesterol levels were normalized to those in the corresponding shCtrl cells expressing an empty vector (Vec.). **(C)** Total cellular cholesterol levels and HMGCR protein expression following site-specific installation of m^6^A on *HMGCR* mRNA using dCas13b-METTL3nls (nuclear localization signal). Cells expressed control sgRNA (sgCtrl) or HMGCR-targeting sgRNA (sgHMGCR) together with wild-type dCas13b-METTL3nls (WT) or the catalytically inactive METTL3 mutant (CD). Cholesterol levels were normalized to those in cells expressing sgCtrl and wild-type dCas13b-METTL3nls. Actin was served as the loading control. Blots are representative of three independent experiments. **(D)** Total cellular cholesterol levels in LNCaP cells expressing wild-type (M3-WT and N93-WT) or mutant forms (M3-CD and N93-AP) of METTL3 and NUP93 under hormone-depleted conditions. Cholesterol levels were normalized to those in cells expressing the corresponding empty vector (Vec.). **(E)** Schematic representation of the steroidogenic pathway through which cholesterol is converted to dihydrotestosterone (DHT). **(F)** Cellular DHT levels in LNCaP cells expressing wild-type (M3-WT and N93-WT) or mutant forms (M3-CD and N93-AP) of METTL3 and NUP93 under hormone-depleted conditions. **(G)** Effects of site-specific removal of m^6^A from *HMGCR* mRNA on HMGCR m^6^A enrichment (left) and cellular DHT levels (right) in LNCaP cells expressing an empty vector (Vec.) or wild-type METTL3 (M3-WT) under hormone-depleted conditions. Cells expressed wild-type dCasRx-ALKBH5nls together with sgCtrl or sgHMGCR. m^6^A enrichment was measured by m^6^A-RIP-qPCR and is presented as the percentage of methylated *HMGCR* RNA relative to input. **(H)** Effects of site-specific removal of m^6^A from *HMGCR* mRNA on HMGCR m^6^A enrichment (left) and cellular DHT levels (right) in LNCaP cells expressing an empty vector (Vec.) or N93-WT under hormone-depleted conditions. Cells expressed wild-type dCasRx-ALKBH5nls together with sgCtrl or sgHMGCR. m^6^A enrichment was measured by m^6^A-RIP–qPCR and is presented as the percentage of methylated *HMGCR* RNA relative to input. Data in (A-D and F-H) are presented as mean ± SD from three independent experiments. *P* values were determined by two-tailed paired Student’s *t* tests for the indicated comparisons.

It is well established that prostate cancer cells sustain AR signaling through intratumoral androgen synthesis following systemic androgen deprivation, thereby promoting tumor growth and the development of CRPC (36,37). Because cholesterol serves as a precursor for intratumoral androgen synthesis (38), our data prompted us to investigate whether overexpression of METTL3 and NUP93 is sufficient to drive castration resistant phenotypes in prostate cancer cells. To this end, we first determined the cholesterol levels in the androgen-dependent prostate cancer cell line LNCaP under hormone depleted condition and found that only when the wild-type METTL3 or NUP93, but not M3-CD nor N93-AP, was overexpressed, nuclear export of mRNAs involved in cholesterol metabolism was markedly enhanced (**Supplemental Fig. 4C-D**), the protein levels of corresponding enzymes were increased (**Supplemental Fig. 4E**), and cellular cholesterol levels was significantly elevated (**Fig. 4D**).

We further sought to confirm that it is the *de novo* synthesis rather than the uptake or efflux process that causes increase in cholesterol levels by measuring cholesterol internalization using fluorescent NBD-cholesterol. Overexpression of wild-type or mutant forms of METTL3 and NUP93 did not change the take-up signals of NBD-cholesterol, indicating that the elevated cholesterol levels were not due to enhanced uptake capacity (**Supplemental Fig. 4F**). These findings support a model in which overexpression of METTL3 and NUP93 augments intracellular cholesterol primarily through m^6^A-dependent nuclear export of transcripts encoding the biosynthesis enzymes rather than through changes in cholesterol import.

Finally, we examined the effects of m^6^A-METTL3-NUP93 axis on intracellular androgen levels in prostate cancer cells under castrate condition (**Fig. 4E**). Overexpression of the wild-type METTL3 or NUP93 increased dihydrotestosterone (DHT) levels in LNCaP cells even under hormone-depleted condition (**Fig. 4F**), whereas M3-CD and N93-AP failed to do so. Importantly, site-specific removal of m^6^A marks on *HMGCR* mRNAs using two separate CRISPR-based tools (dCasRx-ALKBH5nls and dCas13b-ALKBH5nls) attenuated the increases in cellular cholesterol, HMGCR protein abundance, and DHT levels, which was driven by METTL3 or NUP93 overexpression (**Fig. 4G-H** and **Supplemental Fig. 5**). These results directly link the efficient nuclear export mediated by the m^6^A-METTL3-NUP93 axis to intracellular steroidogenesis and androgen biosynthesis in prostate cancer cells under castrate condition.

### METTL3-NUP93 axis integrates metabolic rewiring with androgen receptor reactivation in CRPC

The next question is obviously whether the m^6^A-METTL3-NUP93 axis activates the androgen receptor (AR) under castrate condition. Using a luciferase reporter containing *KLK3* enhancer elements, overexpression of either wild-type METTL3 or NUP93 robustly enhanced the luciferase activity as strong as the addition of 1-2 nM DHT, suggesting the activation of AR signaling in the absence of exogenous ligand (**Fig. 5A**).We also performed AR ChIP-seq (**Fig. 5B**), and found that wild-type METTL3 or NUP93, but not their mutant forms (M3-CD and N93-AP), induced chromatin recruitment of AR at DNA sequences displaying canonical androgen response elements (AREs) and consensus forkhead DNA motifs (**Fig. 5C**). Indeed, at the promoter and enhancer regions of classical DHT-induced genes like *KLK2*, *KLK3*, and *TMPRSS2*, there was dominant accumulation of AR binding (**Fig. 5D** and **Supplemental Fig. 6A-B**). In agreement, gene expression profiling revealed that overexpression of wild-type METTL3 or NUP93, but not the catalytically dead METTL3 or METTL3-binding-defective NUP93 mutant, activated the same group of genes in LNCaP cells under androgen depleted conditions (**Fig. 5E**), which are enriched for androgen response, indicating the activation of androgen-AR signaling (**Fig. 5F-G**). Across CRPC cohorts (39), METTL3 and NUP93 expression both showed a significantly positive correlation with an AR activity score (**Fig. 5H**). All the evidence demonstrate that the m^6^A-METTL3-NUP93 axis reinforces castration-resistant states by activating AR signaling through intracellular androgen production from cholesterol.

**Figure 5.**
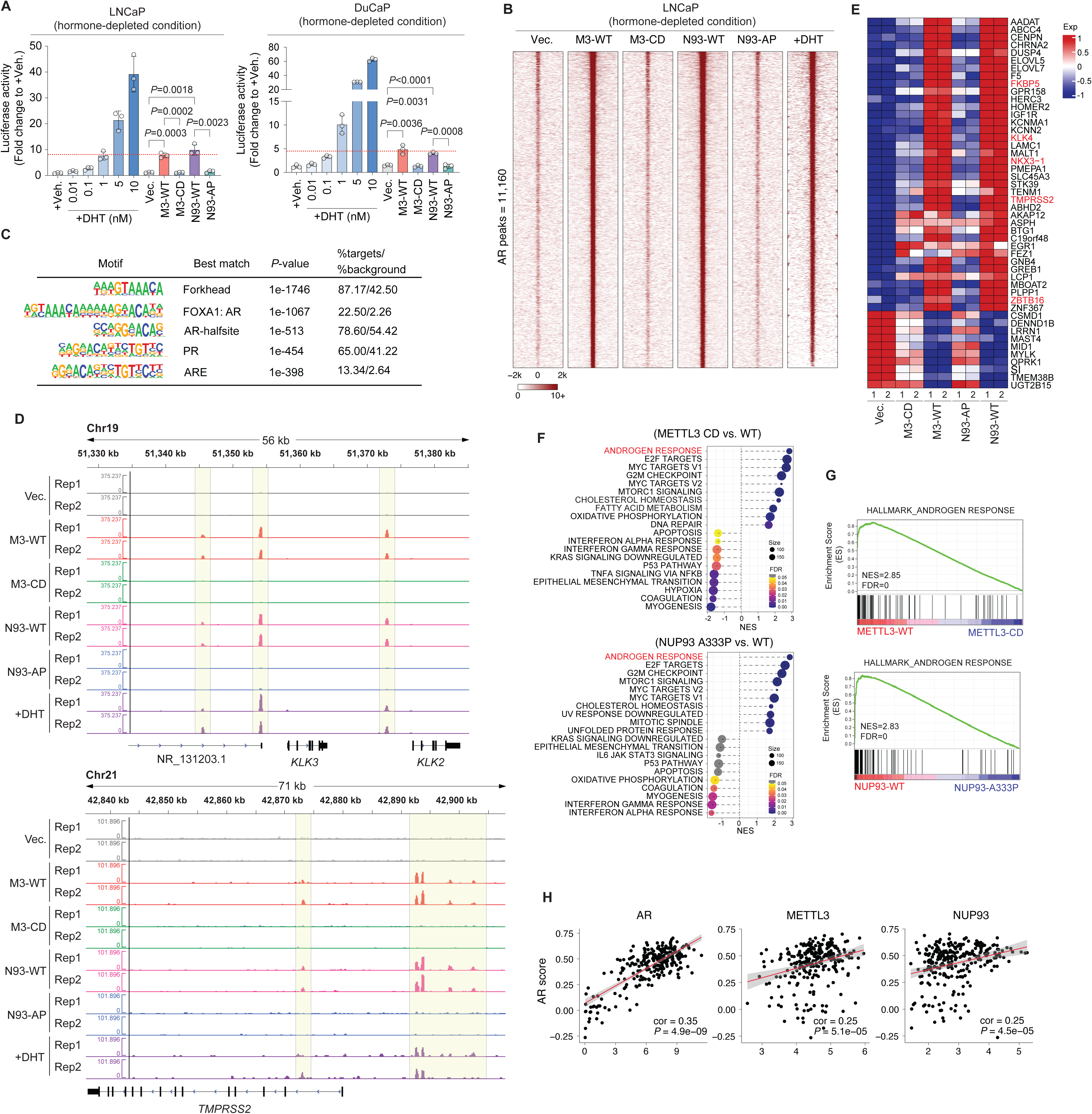
Cholesterol biosynthesis induced by m^6^A-METTL3-NUP93 axis reactivates androgen receptor signaling in the absence of exogenous androgen. **(A)** Androgen receptor (AR) transcriptional activity in LNCaP (left) and DuCaP (right) cells expressing wild-type METTL3 (M3-WT), catalytically inactive METTL3 (M3-CD), wild-type NUP93 (N93-WT), or the METTL3-binding-defective NUP93 mutant (N93-AP) under hormone-depleted conditions. Cells were treated with the indicated concentrations of dihydrotestosterone (DHT) for 16 hrs following hormone depletion as positive controls. AR activity was measured using a *KLK3* enhancer-driven luciferase reporter and is presented relative to that in vehicle (ethanol)-treated cells (+Veh.). **(B)** Heatmaps showing AR ChIP-seq signals across 11,160 AR-binding sites in LNCaP cells expressing an empty vector (Vec.), wild-type METTL3 (M3-WT), catalytically dead METTL3 (M3-CD), wild-type NUP93 (N93-WT), or METTL3-binding-deficient NUP93 (N93-AP) under hormone-depleted conditions. Cells were treated with 2 nM DHT for 16 hrs following hormone depletion as a positive control. Signals are shown within 2 kb upstream and downstream of the AR peak centers. **(C)** Enriched transcription factor-binding motifs identified within the AR-binding sites shown in (B). The best-matched motif, enrichment *P* value, and percentage of target and background sequences containing each motif are shown. **(D)** Genome-browser tracks showing AR ChIP-seq signals from two biological replicates (Rep1 and Rep2) at the *KLK3/KLK2* (top) and *TMPRSS2* (bottom) loci in LNCaP cells expressing an empty vector (Vec.), wild-type METTL3 (M3-WT), catalytically dead METTL3 (M3-CD), wild-type NUP93 (N93-WT), or METTL3-binding-deficient NUP93 (N93-AP) under hormone-depleted conditions. Cells treated with 2 nM DHT (+DHT) were included as a positive control. Highlighted regions indicate AR-binding sites. **(E)** Heatmap showing the expression of androgen-responsive genes in LNCaP cells expressing an empty vector, M3-CD, M3-WT, N93-AP, or N93-WT under hormone-depleted conditions. Two biological replicates are shown for each condition. Selected canonical AR target genes are highlighted in red. **(F)** Hallmark gene-set enrichment analysis (GSEA) comparing LNCaP cells expressing wild-type METTL3 (M3-WT) with those expressing catalytically dead METTL3 (M3-CD) (top) and cells expressing wild-type NUP93 (N93-WT) with those expressing METTL3-binding-defective NUP93 (N93-AP) (bottom). Circle position and color represent the normalized enrichment score (NES), circle size represents the number of genes in each gene set, and *FDR* values are indicated by the color scale. The androgen-response pathway is highlighted in red. **(G)** Gene-set enrichment running score plots showing enrichment of the Androgen Response gene set (Hallmark) in cells expressing wild-type METTL3 (M3-WT) relative to those expressing catalytically dead METTL3 (M3-CD) (top) and in cells expressing wild-type NUP93 (N93-WT) relative to those expressing METTL3-binding-defective NUP93 (N93-AP) (bottom). NES and *FDR* values are shown. **(H)** Correlation of the AR activity score with AR (left), METTL3 (middle), or NUP93 (right) expression in the SU2C dataset (62). Correlation coefficients and *P* values are shown.

### METTL3-NUP93 axis–dependent cholesterol/DHT production rewires LNCaP cells toward a CRPC-like state

At last, we wanted to demonstrate that overexpression of METTL3 and NUP93 induces the castration-resistant phenotypes in prostate cancer cells. Indeed, overexpression of wild-type METTL3 or NUP93, but not their mutant forms, enabled hormone-independent growth (**Fig. 6A** and **Supplemental Fig. 7A-B**), anchorage-independent colony formation (**Fig. 6B**), and enhanced migration and invasion (**Fig. 6C-D**) of androgen-dependent prostate cancer cells under hormone depleted conditions, showing the characteristics of castration-resistant progression.

**Figure 6.**
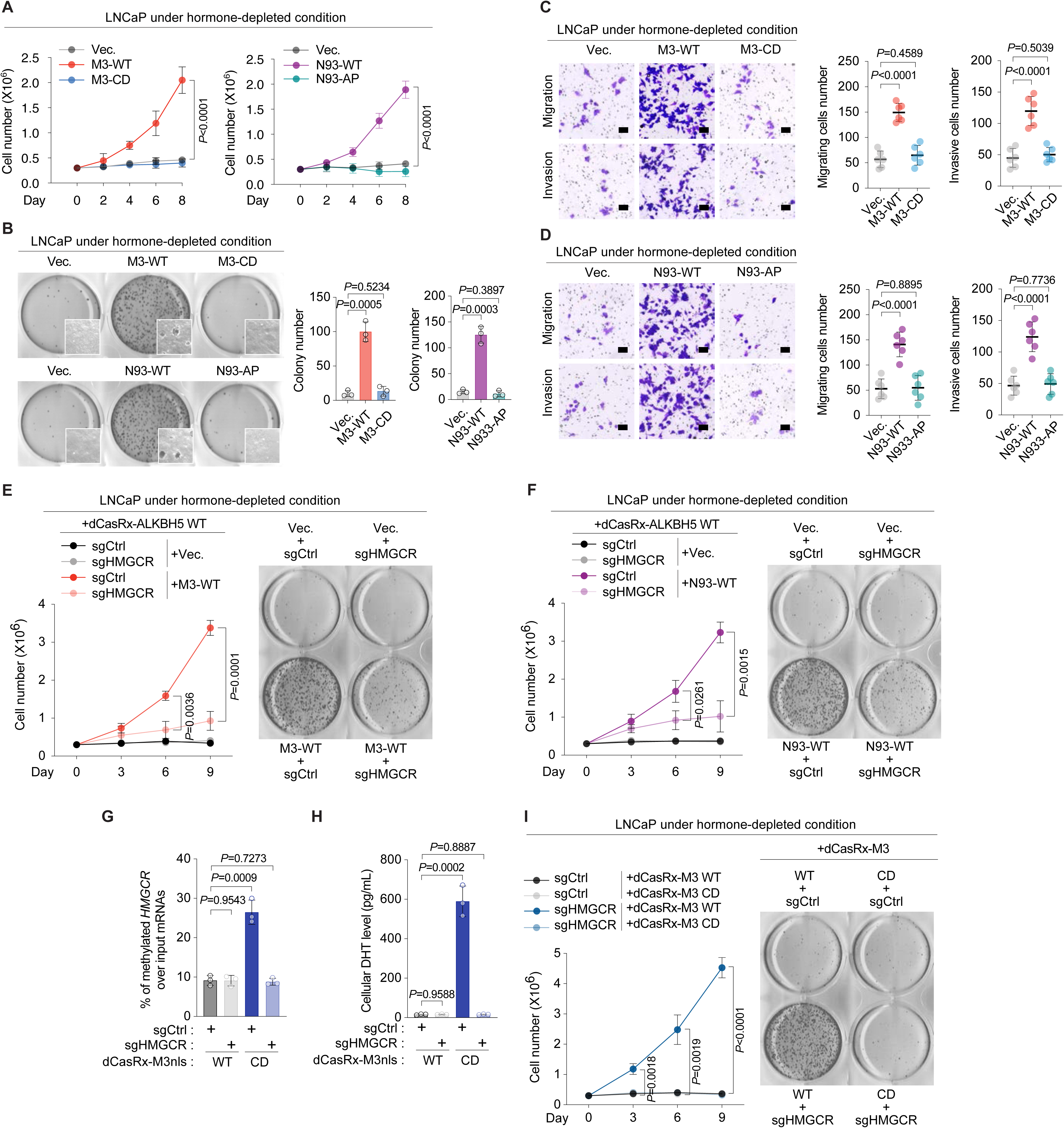
Site-specific m^6^A modification of *HMGCR* mRNA controls acquisition of castration-resistant phenotypes. **(A)** Proliferation of LNCaP cells expressing an empty vector (Vec.), wild-type METTL3 (M3-WT), catalytically inactive METTL3 (M3-CD), wild-type NUP93 (N93-WT), or the METTL3-binding-defective NUP93 mutant (N93-AP) under hormone-depleted conditions. **(B)** Anchorage-independent colony formation by LNCaP cells expressing an empty vector (Vec.), M3-WT, M3-CD, N93-WT, or N93-AP under hormone-depleted conditions. Representative images are shown on the left, with the corresponding quantification shown on the right. **(C, D)** In vitro transwell migration and invasion assays in LNCaP cells expressing an empty vector (Vec.), M3-WT, or M3-CD (C) or an empty vector (Vec.), N93-WT, or N93-AP (D) under hormone-depleted conditions. Representative images of migrating cells (top) and invading cells (bottom) are shown on the left, with the corresponding quantification shown on the right. Scale bars, 100 μm. **(E)** Effects of site-specific removal of m^6^A from *HMGCR* mRNA on proliferation and anchorage-independent colony formation in LNCaP cells expressing an empty vector (Vec.) or wild-type METTL3 (M3-WT) under hormone-depleted conditions. Cells expressed wild-type dCasRx-ALKBH5nls (nuclear localization signal) together with control sgRNA (sgCtrl) or HMGCR-targeting sgRNA (sgHMGCR). Proliferation curves are shown on the left, and representative images of anchorage-independent colonies are shown on the right. **(F)** Effects of site-specific removal of m^6^A from *HMGCR* mRNA on proliferation and anchorage-independent colony formation in LNCaP cells expressing an empty vector (Vec.) or wild-type NUP93 (N93-WT) under hormone-depleted conditions. Cells expressed wild-type dCasRx-ALKBH5nls together with control sgRNA (sgCtrl) or HMGCR-targeting sgRNA (sgHMGCR). Proliferation curves are shown on the left, and representative images of anchorage-independent colonies are shown on the right. **(G, H)** Site-specific installation of m^6^A on *HMGCR* mRNA and its effect on cellular DHT production in LNCaP cells under hormone-depleted conditions. Cells expressed wild-type dCasRx-METTL3nls (dCasRx-M3nls-WT) or the catalytically inactive mutant (dCasRx-M3nls-CD) together with sgCtrl or sgHMGCR. m^6^A enrichment on *HMGCR* mRNA was measured by m^6^A-RIP-qPCR and is presented as the percentage of methylated *HMGCR* RNA relative to input (G). Cellular DHT levels are shown in (H). **(I)** Effects of site-specific installation of m^6^A on *HMGCR* mRNA on LNCaP cell proliferation and anchorage-independent colony formation under hormone-depleted conditions. Cells expressed wild-type dCasRx-METTL3nls or the catalytically inactive mutant together with sgCtrl or sgHMGCR. Proliferation curves are shown on the left, and representative images of anchorage-independent colonies are shown on the right. Data in (A-I) are presented as mean ± SD from three independent experiments. *P* values were determined by two-tailed paired Student’s *t* tests for the indicated comparisons. Images are representative of three independent experiments.

To determine whether the CRPC phenotypes induced upon overexpression of METTL3 or NUP93 requires the m^6^A modification on their target mRNAs, we utilized the dCasRx-ALKBH5 system to remove site-specific m^6^A marks on *HMGCR* mRNAs. Strikingly, delivery of the target sgRNA fully suppressed hormone-independent growth and anchorage-independent colony formation conferred upon METTL3 or NUP93 overexpression (**Fig. 6E-F**). Conversely, targeted installation of methylation on *HMGCR* mRNAs using dCas13b-METTL3nls tool was sufficient to induce hormone-independent proliferation and anchorage-independent growth of LNCaP cells co-expressing the target sgRNA and wild-type METTL3 compared to those expressing control sgRNA or enzymatically inefficient METTL3 mutant, which is accompanied by the increased intracellular DHT levels (**Fig. 6G-I**). All these results highlight the essential role of m^6^A modification on the target mRNAs of the METTL3-NUP93 complex in driving CRPC progression. Neither AR-V7 expression nor AKT pathway was induced following METTL3 or NUP93 overexpression (**Supplemental Fig. 7C-F**). Thus, CRPC phenotypes driven by the m^6^A-METTL3-NUP93 axis are unlikely the secondary effects of other reported mechanisms leading to CRPC progression.

Taken together, our findings establish the m^6^A-METTL3-NUP93 axis as a novel driver of CRPC progression via facilitating methylation-dependent nuclear export of mRNAs functioning in cholesterol biosynthesis and subsequently increasing intracellular androgen amounts and activating AR signaling under castrate condition.

### Pharmacologic disruption of the m^6^A–METTL3–NUP93 axis suppresses castration-resistant prostate cancer

We next used pharmacological interventions to determine whether the castration-resistant phenotypes driven by the m^6^A-METTL3-NUP93 axis depend on cholesterol biosynthesis, intratumoral androgen production, and downstream AR signaling **(Fig. 7A**). Inhibition of HMGCR and squalene epoxidase (SQLE) with their specific small-molecule inhibitors simvastatin (40) and terbinafine (41), respectively, significantly impaired the growth of LNCaP and DuCaP cells overexpressing the wild-type METTL3 or NUP93 under hormone depleted conditions (**Fig. 7B-C** and **Supplemental Fig. 8A-B**). Likewise, pharmacological blockade of the androgen-AR signaling axis with the CYP17A1 inhibitor abiraterone (42) or the AR antagonist enzalutamide (43) markedly suppressed the castration-resistant phenotypes induced by METTL3 or NUP93 overexpression in LNCaP and DuCaP cells (**Fig. 7B-C** and **Supplemental Fig. 8A-B**). Consistently, each intervention attenuated the induction of classical AR target genes, including *KLK3*, *KLK2*, *TMPRSS2*, and *FKBP5* (**Supplemental Fig. 8C**). Together, these pharmacological rescue experiments establish that the phenotypes induced by METTL3, or NUP93 overexpression are mediated, at least in part, through cholesterol-dependent androgen production and AR activation.

**Figure 7.**
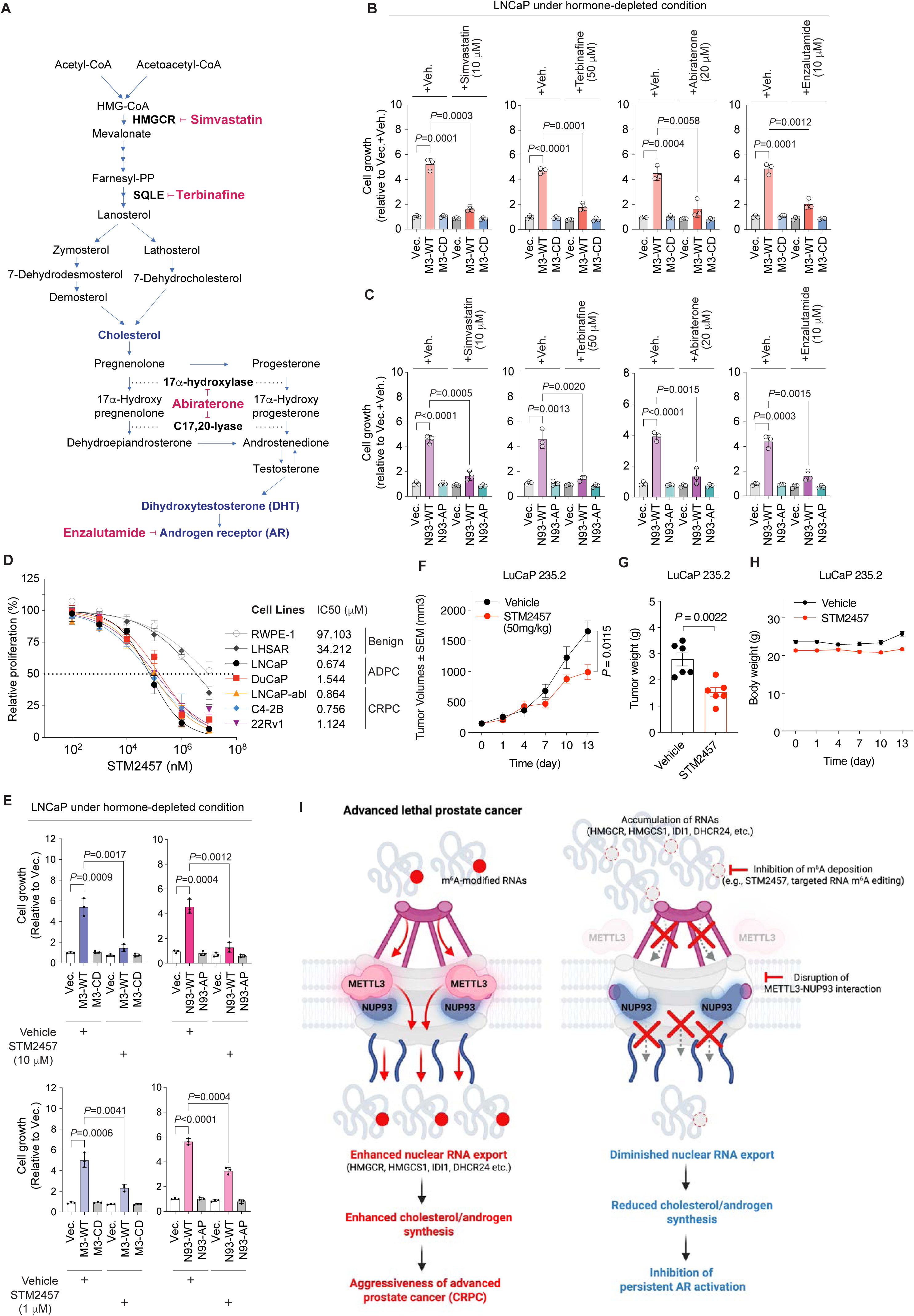
Pharmacologic inhibition of the m^6^A-METTL3-NUP93 axis suppresses castration-resistant prostate cancer. **(A)** Schematic representation of the cholesterol biosynthesis, androgen synthesis, and AR signaling pathways and the sites of action for simvastatin, terbinafine, abiraterone, and enzalutamide. Simvastatin and terbinafine inhibit HMGCR and SQLE, respectively; abiraterone inhibits CYP17A1; and enzalutamide inhibits AR. **(B, C)** Growth of LNCaP cells expressing an empty vector (Vec.), wild-type METTL3 (M3-WT), or catalytically inactive METTL3 (M3-CD) (B) and an empty vector (Vec.), wild-type NUP93 (N93-WT), or the METTL3-binding-defective NUP93 mutant (N93-AP) (C) under hormone-depleted conditions. Cells were treated with vehicle (ethanol, Veh.), simvastatin (10 μM), terbinafine (50 μM), abiraterone (20 μM), or enzalutamide (10 μM) for 5 days. Cell growth was normalized to that of the corresponding vehicle-treated vector control (Vec.). **(D)** Dose-response analysis of METTL3 inhibitor STM2457 following 72 hrs treatment in benign prostate epithelial cells (RWPE-1 and LHSAR), androgen-dependent prostate cancer cells (LNCaP and DuCaP), and CRPC cells (LNCaP-abl, C4-2B, and 22Rv1). Relative proliferation and calculated IC_50_ values are shown. **(E)** Cell growth of LNCaP cells expressing an empty vector (Vec.), M3-WT, M3-CD, N93-WT, or N93-AP under hormone-depleted conditions following treatment with vehicle or STM2457 at 10 μM (top) or 1 μM (bottom) for 5 days. Cell growth was normalized to that of the corresponding vehicle-treated vector control (Vec.). **(F-H)** Therapeutic effects of STM2457 in the patient-derived xenograft model of CRPC (LuCaP 235.2). Mice were treated with vehicle or STM2457 (50 mg/kg). Tumor growth over the indicated period (F), endpoint tumor weights (G), and changes in mouse body weight during treatment (H) are shown. **(I)** Proposed model depicting the role of the m^6^A-METTL3-NUP93 axis in advanced prostate cancer. Deposition of m^6^A modification and METTL3-NUP93 cooperation promote the nuclear export of transcripts encoding cholesterol biosynthesis enzymes, thereby increasing cholesterol and androgen synthesis and promoting AR activation and CRPC progression. Inhibition of m^6^A deposition or disruption of the METTL3-NUP93 interaction impairs nuclear RNA export, reduces cholesterol and androgen synthesis, suppresses persistent AR activation, and inhibits prostate cancer progression. Quantitative data in (B-E) are presented as mean ± SD from three independent experiments. *P* values were determined by two-tailed paired Student’s *t* tests for the indicated comparisons. Data in (F-H) are presented as mean ± SEM. *P* values in (F) and (G) were determined by two-tailed unpaired Student’s *t* tests.

We next asked whether pharmacological disruption of the m^6^A-METTL3-NUP93 axis itself using the small-molecule inhibitor of METTL3, STM2457, is effective in treating prostate cancer, especially CRPC. Dose-response analysis across multiple prostate epithelial cell lines, representing benign, androgen-dependent, and castration-resistant models, revealed clearly more sensitivity to STM2457 in prostate adenocarcinoma cell lines (with IC_50_ values below 2 μM in general) than in the benign ones (with IC_50_ values more than 30 μM) (**Fig. 7D**). This substantial therapeutic window underscores the feasibility of targeting aberrant signaling that involves m^6^A modification for the treatment of prostate cancer. STM2457 treatment markedly inhibited the CRPC phenotypes that are induced upon overexpression of METTL3 or NUP93 in androgen-dependent prostate cancer cells (**Fig. 7E** and **Supplemental Fig. 8D**). Interestingly, STM2457 treatment also suppressed the upregulation of androgen-responsive genes like *KLK3* and *TMPRSS2*, only when cells overexpressed the wild-type METTL3 or NUP93 (**Supplemental Fig. 8E**). In two independent CRPC cell lines, C4-2B and 22Rv1, STM2457 treatment reduced m^6^A levels on transcripts encoding cholesterol biosynthetic enzymes (*HMGCR*, *HMGCS1*, *IDI1*, *DHCR24*) and promoted their nuclear retention without altering their total abundance (**Supplemental Fig. 8F-I**). These effects strongly indicated that the mechanism of action of STM2457 in prostate cancer is likely to impair m^6^A-dependent nuclear export of transcripts functionally involved in cholesterol metabolism and further to block intratumoral activation of androgen-AR signaling. Finally, in the patient-derived xenograft model of CRPC (LuCaP 235.2), STM2457 treatment significantly retarded tumor growth without affecting animal body weight (**Fig. 7F-H**).

Collectively, our findings identify the m^6^A-METTL3-NUP93 axis as a therapeutically targetable driver of CRPC progression. Pharmacologic inhibition of METTL3 disrupts this axis, suppresses cholesterol-dependent reactivation of AR signaling, and inhibits CRPC growth.

## Discussion

Previous work, including ours, has demonstrated that NPCs and m^6^A machinery work together to facilitate the nuclear export of selective mRNAs (23), which regulate a variety of physiological processes (44). However, the function of their crosstalk in cancer has somehow been neglected. In the present study, we unveil strong oncogenic features of the nucleoporin protein NUP93 in prostate cancer cells, which are dependent on its interaction with METTL3. The METTL3-NUP93 complex expedites the passage of particular mRNA molecules with m^6^A modification through NPCs. This distinct mRNA transport pathway promotes progression of prostate cancer to castration resistance, revealing an important function of m^6^A-NPC cooperation in advanced prostate cancer. We found that direct mRNA cargoes of the m^6^A-METTL3-NUP93 axis are functionally enriched in the biosynthesis of cholesterol, the natural precursor for androgen synthesis (45). Thus, overexpression of the wild-type METTL3 or NUP93, but not their mutant forms that impede the mRNA transport axis, elevated the cellular DHT levels, induced AR activation under castrate condition, conferred androgen-independent growth, and promoted migration and invasion of hormone sensitive prostate cancer cells. All these results indicate the emergence of castration-resistant prostate cancer. This study provides the original evidence showing that the cooperation between NPCs and m^6^A machinery plays an important role in prostate cancer progression, promoting the aggressive transformation of the disease from androgen dependent to androgen independent and thereby holding promise as new therapeutic targets for treating advanced and castration-resistant prostate cancer (**Fig. 7I**).

Although NPCs have been shown to be vulnerable in several cancers, directly targeting these essential cellular structures for cancer intervention is generally considered unfeasible and toxic. Previously, it was demonstrated that the nucleoporin protein POM121 promotes lethal prostate cancer by facilitating importin β-dependent nuclear import of key oncogenic transcription factor such as E2F1 and MYC (35). Therefore, treatment with importin β inhibitor decreased tumor growth in xenograft models of advanced prostate cancer. In another study, XPO1/CRM1, the major transport receptor for exporting proteins and multiple RNA species (46,47), was found elevated in metastatic CRPC that has become resistant to AR-targeted therapies (48). XPO1 inhibitor showed antitumor activity in these aggressive CRPC cells both *in vivo* and *in vitro* (49). All these studies demonstrate the potential of targeting NPCs-mediated nucleocytoplasmic transport for cancer therapy, not via directly impeding nucleoporins but rather NPC-associated factors. Here, we discovered that m^6^A machinery cooperates with NPCs to enhance CRPC development, providing a new direction to interfere with NPCs’ functions in cancers. Furthermore, the NPC-m^6^A cooperation may also play an oncogenic role in other cancer types besides prostate cancer, suggesting that blocking m^6^A modification may represent a prevalent way to hinder dysfunction of NPCs in cancers.

In prostate cancer cells, we identified a group of mRNAs which depend on the m^6^A-METTL3-NUP93 axis for their movement out of nucleus. Interestingly, a subset of these mRNA molecules is functionally involved in biosynthesis of cholesterol. High circulating cholesterol levels have been linked to the development of aggressive prostate cancer in clinics (50,51). Previous theories attribute the abnormal cholesterol accumulation to upregulation of lipid uptake from the bloodstream (52,53), downregulation of cholesterol efflux (54,55) and increased synthesis rates (56,57). Here we provide a new mechanism for dysregulated cholesterol homeostasis in cancer, which is caused by expedited nuclear export of mRNAs encoding the major regulatory enzymes involved in cholesterol biosynthesis, such as *HMGCR* and *HMGCS1*. Notably, some of these protein enzymes have long been reported to be significantly overexpressed in prostate cancer compared to adjacent normal tissues, which is associated with poor prognosis of the disease (29,58). Now our findings reveal an answer to the enigmatic question of their misregulation in cancer and suggest an application of the m^6^A signals on these mRNAs as biomarkers indicative of prostate cancer progression, which warrants further validation and quantification in patient cohorts. Additionally, as essential components of lipid rafts, cholesterol and its biosynthesis intermediates have been shown to activate several signaling pathways that drive malignancies, such as PI3K/AKT/mTOR, Hedgehog signaling, and MAPK signaling cascades, etc (59). Although our studies showed no changes in the active forms of AKT when the wild-type or mutant forms of METTL3 and NUP93 were overexpressed, we cannot exclude the possibility that activation of other oncogenic signaling pathways contributes to the promotion of CRPC development mediated by the m^6^A-METTL3-NUP93 axis. Nevertheless, our work highlights the potentials of this new mRNA export pathway as an anticancer target and biomarkers for advanced prostate cancer.

From a therapeutic perspective, our pharmacological rescue experiments established that cholesterol-dependent activation of AR signaling is the downstream effector mediating METTL3- or NUP93-driven phenotypes. Therefore, pharmacological inhibition of the upstream m^6^A-METTL3-NUP93 axis may be complementary to the conventional AR-directed therapies for advanced prostate cancer. Consistent with this speculation, METTL3 inhibitor STM2457 preferentially inhibited the proliferation of prostate cancer cells, including CRPC lines, and suppressed the tumor growth in the PDX model of CRPC without affecting animal body weight. These findings provide proof of concept for targeting the m^6^A-METTL3-NUP93 axis in CRPC. Further studies are warranted to evaluate the potency, selectivity, and safety of METTL3-targeting compounds and to determine whether expression of METTL3 or NUP93 can serve as biomarkers for predicting therapeutic benefits. Furthermore, considering that resistance to AR pathway inhibitors represents a major clinical challenge (60,61), it is appealing to investigate whether pharmacological disruption of the m^6^A-METTL3-NUP93 axis inhibition is effective in overcoming the therapeutic resistance or combination of METTL3-targeting compounds and metabolic enzyme inhibitors or AR antagonists has a synergistic or additive effect in CRPC.

In summary, we demonstrate that the mRNA export mediated by the m^6^A-METTL3-NUP93 axis promotes prostate cancer progression. Mechanistically, we identified a group of mRNAs encoding cholesterol synthesis enzymes, which rely on the m^6^A modification and the METTL3-NUP93 interaction for moving through NPCs to stimulate cholesterol biosynthesis and steroidogenesis, activate the AR signaling axis under castrate condition, and consequently induce castration-resistant phenotypes. Our findings reveal the mechanisms underlying the oncogenic function of NPC-m^6^A crosstalk, as well as lay the foundation for the development and expansion of therapeutic modalities for the treatment of CRPCs and potentially other types of aggressive tumors.

---

**Supplemental Figure S1.**
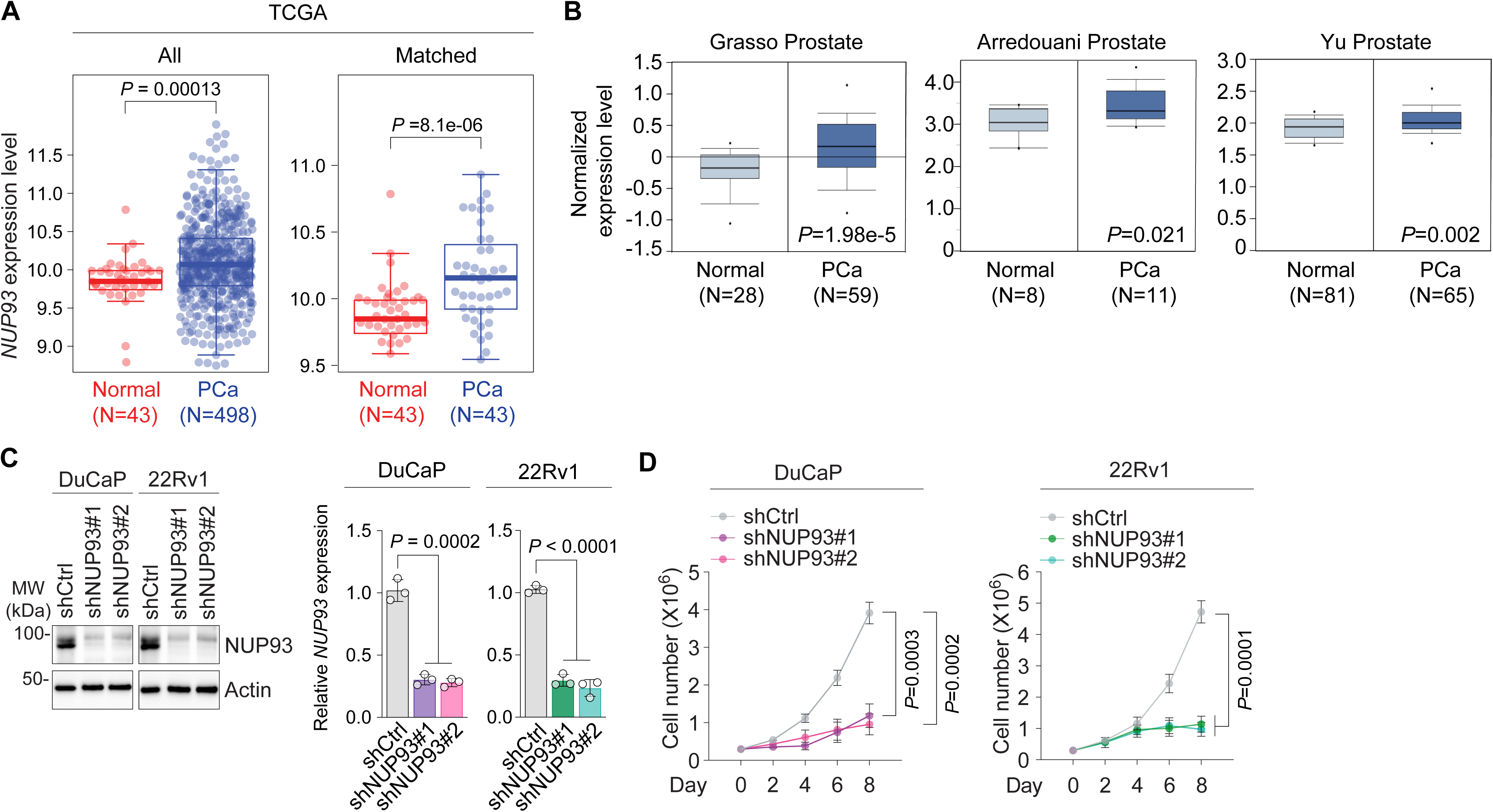
NUP93 is upregulated in prostate cancer and required for prostate cancer cell proliferation. **(A)** *NUP93* expression in nonmalignant prostate tissues and primary prostate cancer (PCa) specimens from TCGA-PRAD. The analysis includes all available nonmalignant (n = 43) and primary tumor (n = 498) samples (left) and 43 matched pairs of nonmalignant and primary tumor tissues (right). *P* values were determined by Wilcoxon rank sum and signed rank test, respectively. **(B)** *NUP93* expression in nonmalignant prostate tissues and primary prostate cancer specimens from the Grasso, Arredouani, and Yu prostate cancer cohorts (GSE35598, GSE55945, and GSE6919). Sample numbers and *P* values are indicated. **(C)** Validation of NUP93 depletion in DuCaP and 22Rv1 cells expressing control shRNA (shCtrl) or NUP93-targeting shRNAs (shNUP93#1 and shNUP93#2). NUP93 protein and mRNA levels were assessed by immunoblotting (left) and RT-qPCR (right), respectively. Actin was served as the loading control for immunoblotting. *NUP93* mRNA expression was normalized to *GAPDH* and presented relative to that in the corresponding shCtrl cells. Blots are representative of three independent experiments. **(D)** Proliferation of DuCaP (left) and 22Rv1 (right) cells expressing shCtrl, shNUP93#1, or shNUP93#2. Quantitative data in (C, D) are presented as mean ± SD from three independent experiments. *P* values were determined by two-tailed paired Student’s *t* tests for the indicated comparisons.

**Supplemental Figure S2.**
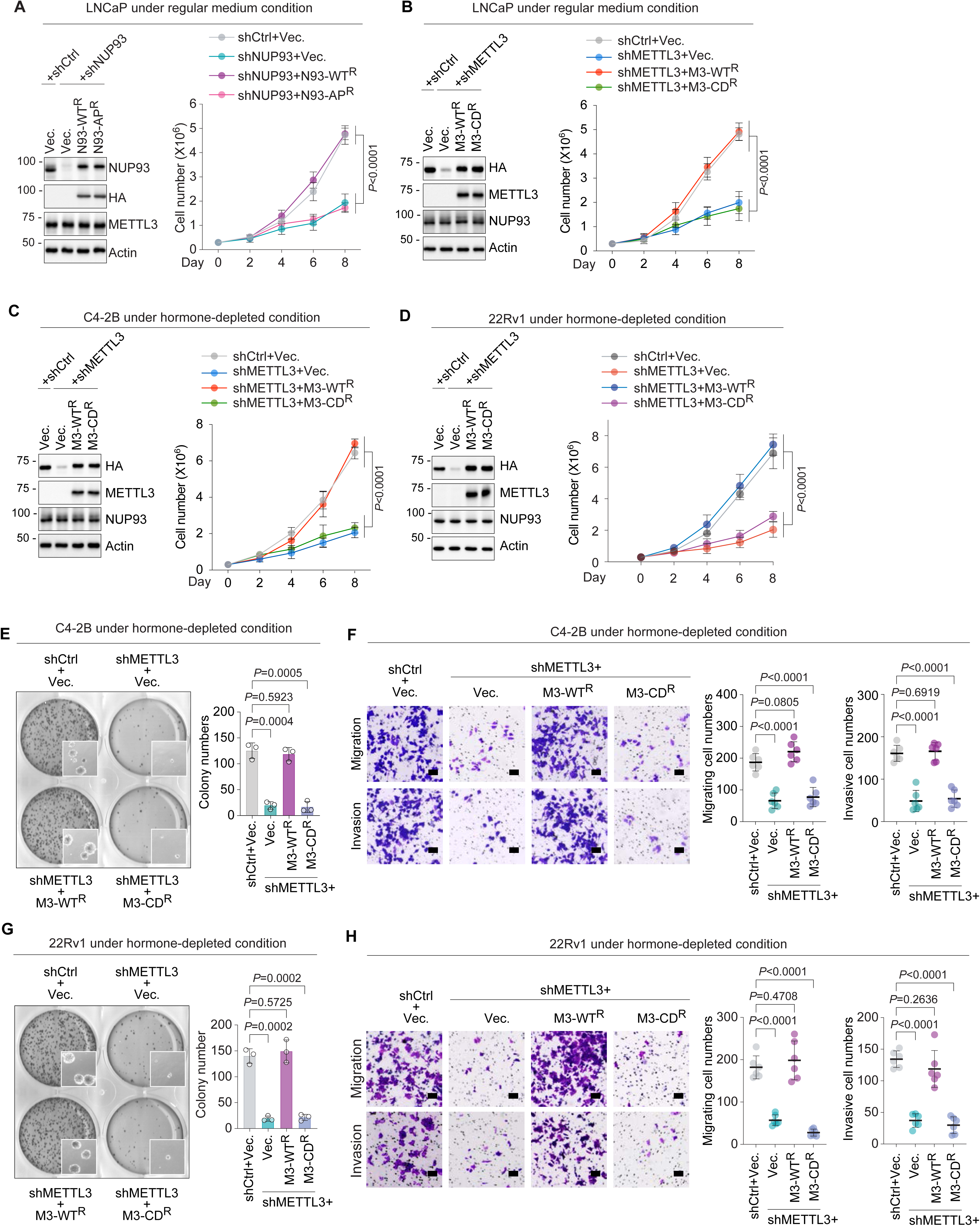
METTL3 catalytic activity and METTL3-NUP93 interaction are required for prostate cancer cell proliferation and malignant phenotypes. **(A)** Rescue of LNCaP cell proliferation following endogenous NUP93 depletion and re-expression of shRNA-resistant wild-type NUP93 (N93-WT^R^) or the METTL3-binding-defective NUP93 mutant (N93-AP^R^) under regular medium conditions. Immunoblot analysis of NUP93, HA-tagged NUP93, and METTL3 is shown on the left, with Actin serving as the loading control. Cell proliferation is shown on the right. **(B)** Rescue of LNCaP cell proliferation following endogenous METTL3 depletion and re-expression of shRNA-resistant wild-type METTL3 (M3-WT^R^) or catalytically inactive METTL3 (M3-CD^R^) under regular medium conditions. Immunoblot analysis of HA-tagged METTL3, total METTL3, and NUP93 is shown on the left, with Actin serving as the loading control. Cell proliferation is shown on the right. **(C, D)** Rescue of cell proliferation following endogenous METTL3 depletion and re-expression of M3-WT^R^ or M3-CD^R^ in C4-2B (C) and 22Rv1 (D) cells under hormone-depleted conditions. Immunoblot analysis of HA-tagged METTL3, total METTL3, and NUP93 is shown on the left, with Actin serving as the loading control. Cell proliferation is shown on the right. **(E, G)** Anchorage-independent colony formation following endogenous METTL3 depletion and re-expression of M3-WT^R^ or M3-CD^R^ in C4-2B (E) and 22Rv1 (G) cells under hormone-depleted conditions. Representative images are shown on the left, with the corresponding quantification shown on the right. **(F, H)** *In vitro* transwell migration and invasion assays following endogenous METTL3 depletion and re-expression of M3-WT^R^ or M3-CD^R^ in C4-2B (F) and 22Rv1 (H) cells under hormone-depleted conditions. Representative images of migrating cells (top) and invading cells (bottom) are shown on the left, with the corresponding quantification shown on the right. Scale bars, 100 μm. Data in (A-H) are presented as mean ± SD from three independent experiments. *P* values were determined by two-tailed paired Student’s *t* tests for the indicated comparisons. Images and blots are representative of three independent experiments.

**Supplemental Figure S3.**
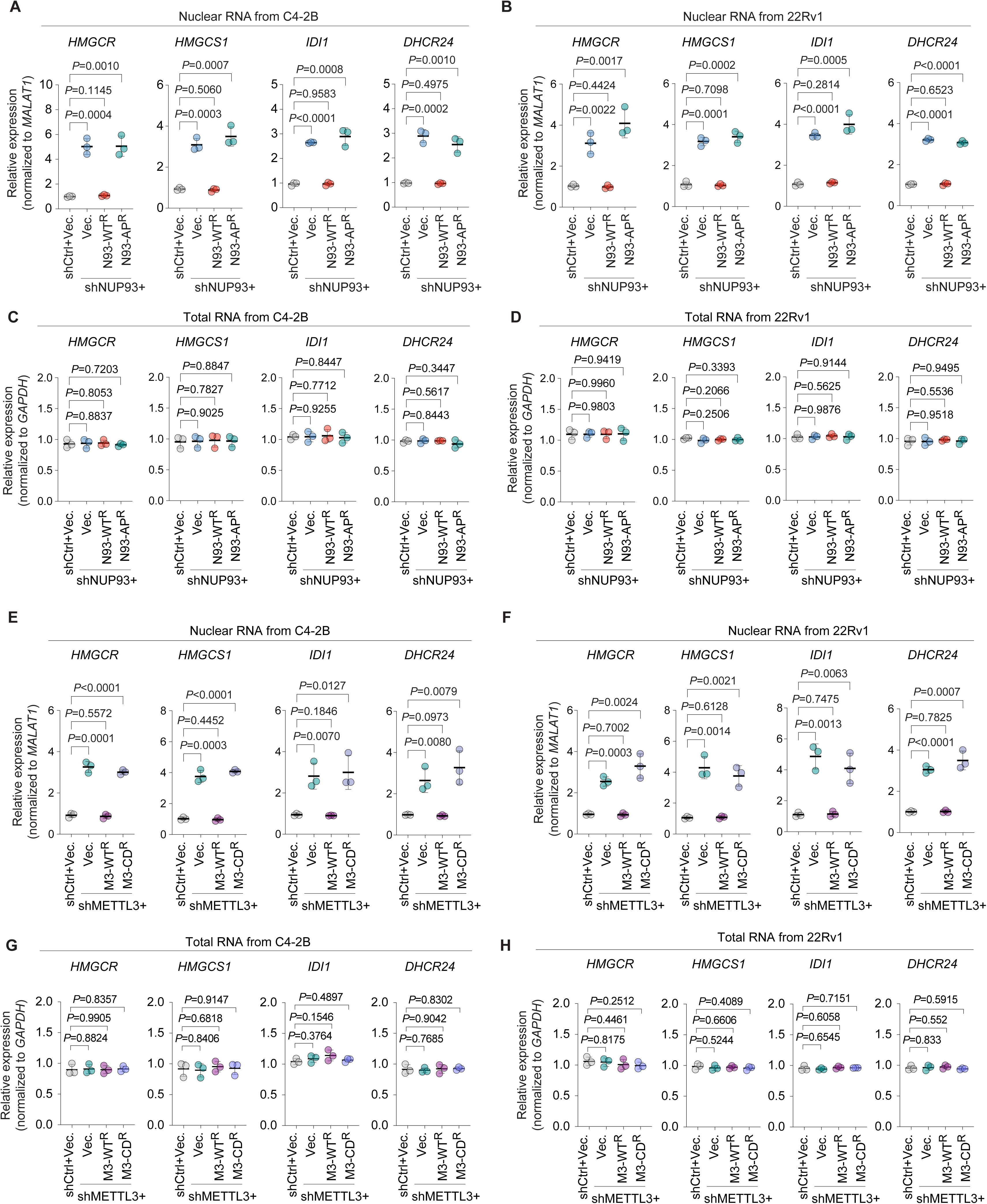
METTL3 catalytic activity and METTL3-NUP93 interaction are required for the nuclear export of cholesterol biosynthesis transcripts. **(A-D)** Nuclear and total RNA levels of *HMGCR*, *HMGCS1*, *IDI1*, and *DHCR24* following endogenous NUP93 depletion and re-expression of shRNA-resistant wild-type NUP93 (N93-WT^R^) or the METTL3-binding-defective NUP93 mutant (N93-AP^R^). Nuclear RNA levels in C4-2B (A) and 22Rv1 (B) cells were normalized to *MALAT1*. Total RNA levels in C4-2B (C) and 22Rv1 (D) cells were normalized to *GAPDH*. Expression levels are presented relative to those in the corresponding shCtrl cells expressing an empty vector (Vec.). **(E-H)** Nuclear and total RNA levels of *HMGCR*, *HMGCS1*, *IDI1*, and *DHCR24* following endogenous METTL3 depletion and re-expression of shRNA-resistant wild-type METTL3 (M3-WT^R^) or catalytically inactive METTL3 (M3-CD^R^). Nuclear RNA levels in C4-2B (E) and 22Rv1 (F) cells were normalized to *MALAT1*. Total RNA levels in C4-2B (G) and 22Rv1 (H) cells were normalized to *GAPDH*. Expression levels are presented relative to those in the corresponding shCtrl cells expressing an empty vector. Data are presented as mean ± SD from three independent experiments. *P* values were determined by two-tailed paired Student’s *t* tests for the indicated comparisons.

**Supplemental Figure S4.**
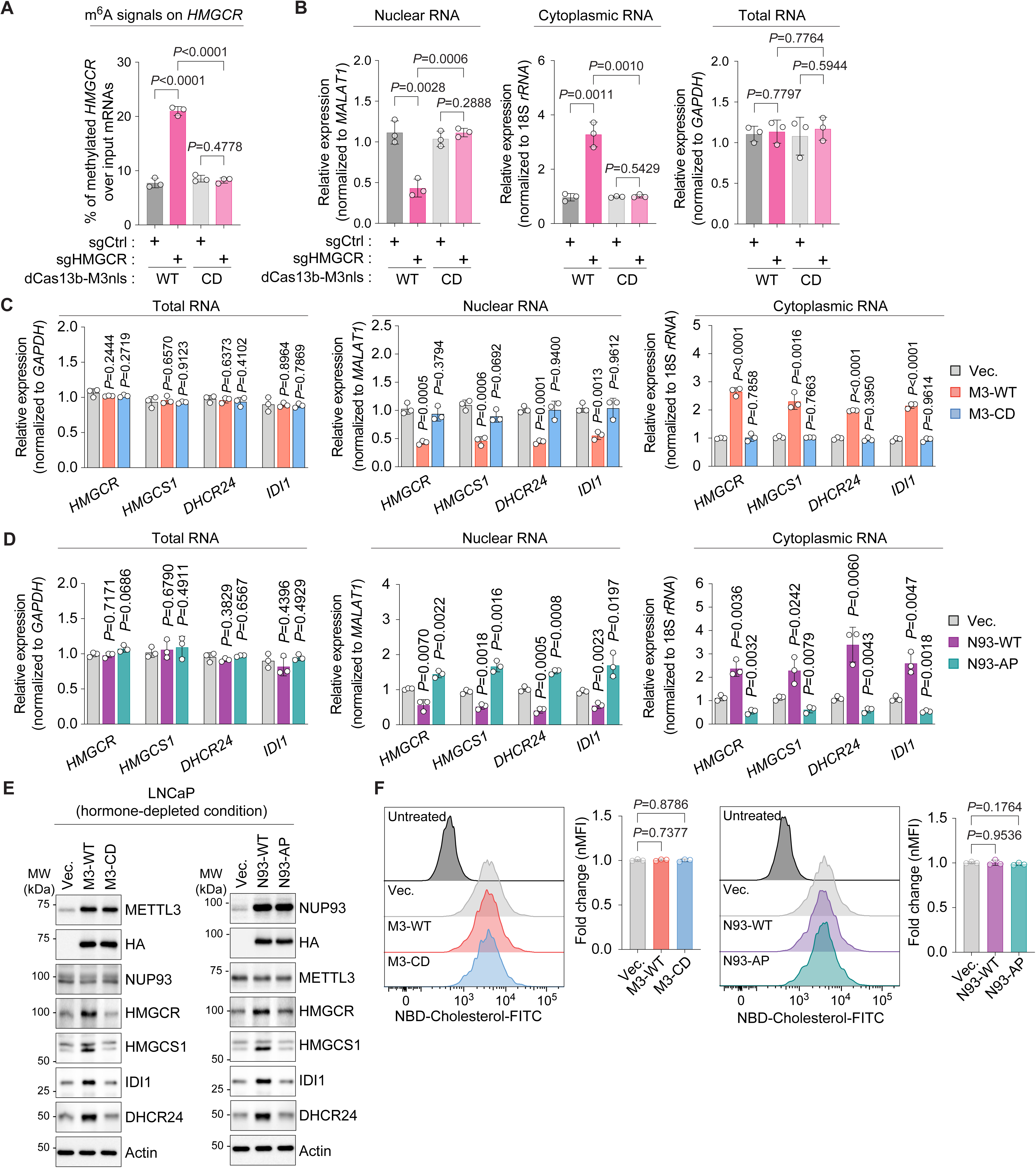
The m^6^A-METTL3-NUP93 axis promotes the nuclear export and expression of cholesterol biosynthesis transcripts without altering cholesterol uptake. **(A)** m^6^A enrichment on *HMGCR* mRNA in cells expressing wild-type dCas13b-METTL3nls (WT) or the catalytically inactive CD mutant together with control sgRNA (sgCtrl) or HMGCR-targeting sgRNA (sgHMGCR). m^6^A enrichment was measured by m^6^A-RIP–qPCR and is presented as the percentage of methylated *HMGCR* RNA relative to input. **(B)** Nuclear (left), cytoplasmic (middle), and total (right) *HMGCR* mRNA levels under the conditions described in (A). Nuclear and cytoplasmic *HMGCR* RNA levels were normalized to *MALAT1* and *18S rRNA*, respectively, and total *HMGCR* RNA was normalized to *GAPDH*. **(C, D)** Total, nuclear, and cytoplasmic RNA levels of *HMGCR*, *HMGCS1*, *DHCR24*, and *IDI1* in LNCaP cells expressing an empty vector (Vec.), M3-WT, or M3-CD (C) or an empty vector (Vec.), N93-WT, or N93-AP (D) under hormone-depleted conditions. Total RNA was normalized to *GAPDH*, whereas nuclear and cytoplasmic RNA levels were normalized to *MALAT1* and *18S rRNA*, respectively. Expression levels are presented relative to those in the corresponding vector-control cells. **(E)** Immunoblot analysis of HA-tagged METTL3 or NUP93, endogenous NUP93 or METTL3, HMGCR, HMGCS1, IDI1, and DHCR24 in LNCaP cells expressing an empty vector (Vec.), M3-WT, M3-CD, N93-WT, or N93-AP under hormone-depleted conditions. Actin was served as the loading control. Blots are representative of three independent experiments. **(F)** NBD-cholesterol uptake in LNCaP cells expressing an empty vector (Vec.), M3-WT, M3-CD, N93-WT, or N93-AP. Representative flow-cytometry histograms and quantification of normalized median fluorescence intensity (nMFI) are shown. Data in (A-D and F) are presented as mean ± SD from three independent experiments. *P* values were determined by two-tailed paired Student’s *t* tests for the indicated comparisons.

**Supplemental Figure S5.**
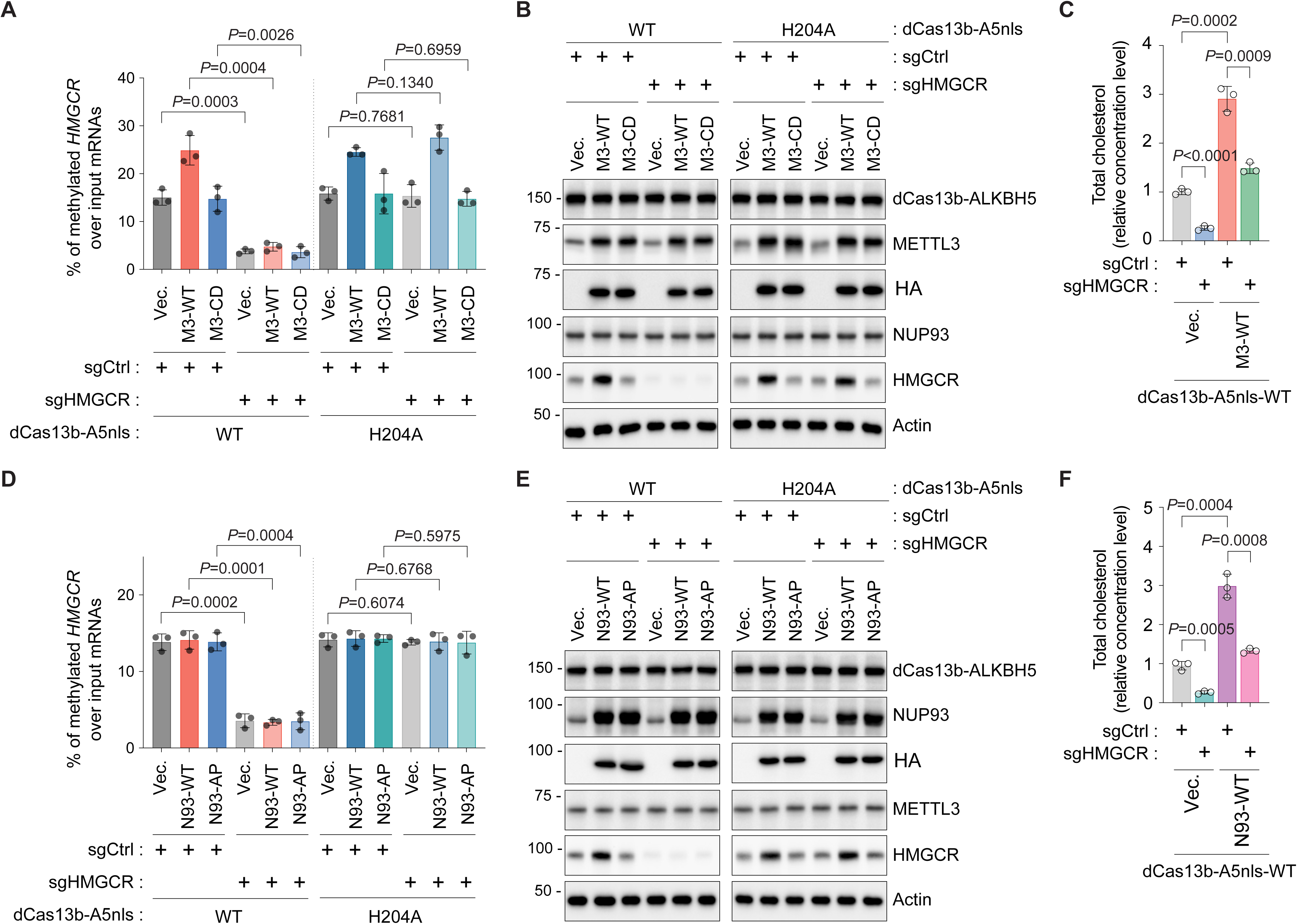
Site-specific removal of m^6^A from HMGCR mRNA attenuates METTL3- and NUP93-induced cholesterol accumulation. **(A)** m^6^A enrichment on *HMGCR* mRNA in LNCaP cells expressing an empty vector (Vec.), wild-type METTL3 (M3-WT), or catalytically inactive METTL3 (M3-CD) under hormone-depleted conditions. Cells expressed wild-type dCas13b-ALKBH5nls (WT) or the catalytically inactive H204A mutant together with control sgRNA (sgCtrl) or HMGCR-targeting sgRNA (sgHMGCR). m^6^A enrichment was measured by m^6^A-RIP-qPCR and is presented as the percentage of methylated *HMGCR* RNA relative to input. **(B)** Immunoblot analysis of dCas13b-ALKBH5nls, METTL3, HA-tagged METTL3, NUP93, and HMGCR under the conditions described in (A). Actin was served as the loading control. Blots are representative of three independent experiments. **(C)** Total cellular cholesterol levels in LNCaP cells expressing an empty vector (Vec.) or M3-WT together with wild-type dCas13b-ALKBH5nls and sgCtrl or sgHMGCR under hormone-depleted conditions. Cholesterol levels were normalized to those in vector-control cells expressing sgCtrl. **(D)** m^6^A enrichment on *HMGCR* mRNA in LNCaP cells expressing an empty vector (Vec.), wild-type NUP93 (N93-WT), or the METTL3-binding-defective NUP93 mutant (N93-AP) under hormone-depleted conditions. Cells expressed wild-type dCas13b-ALKBH5nls or the H204A mutant together with sgCtrl or sgHMGCR. m^6^A enrichment was measured by m^6^A-RIP–qPCR and is presented as the percentage of methylated *HMGCR* RNA relative to input. **(E)** Immunoblot analysis of dCas13b-ALKBH5nls, NUP93, HA-tagged NUP93, METTL3, and HMGCR under the conditions described in (D). Actin was served as the loading control. Blots are representative of three independent experiments. **(F)** Total cellular cholesterol levels in LNCaP cells expressing an empty vector (Vec.) or N93-WT together with wild-type dCas13b-ALKBH5nls and sgCtrl or sgHMGCR under hormone-depleted conditions. Cholesterol levels were normalized to those in vector-control cells expressing sgCtrl. Data in (A, C, D, and F) are presented as mean ± SD from three independent experiments. *P* values were determined by two-tailed paired Student’s *t* tests for the indicated comparisons.

**Supplemental Figure S6.**
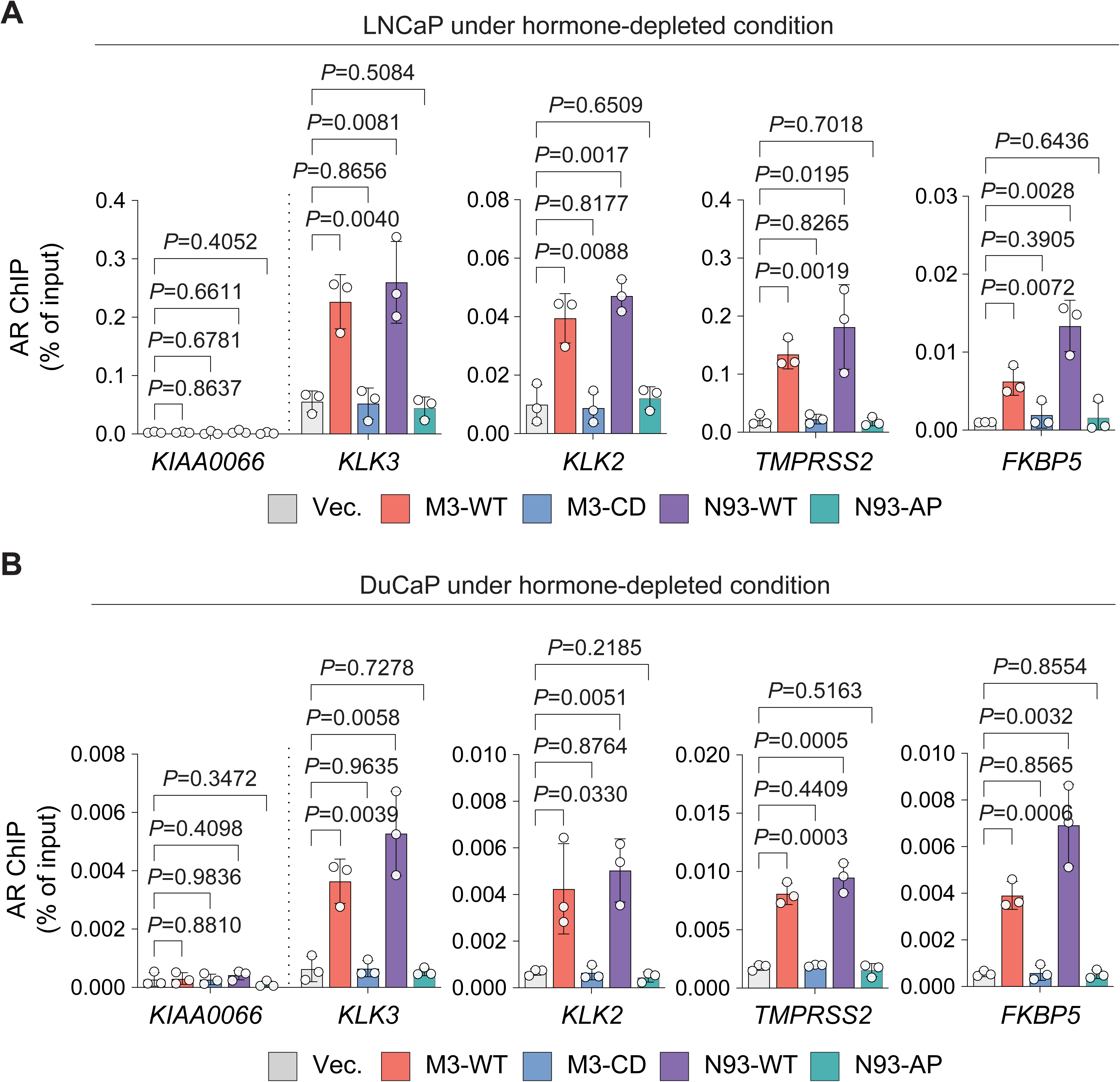
METTL3 catalytic activity and METTL3-NUP93 interaction promote androgen receptor recruitment to AR target genes. **(A-B)** Targeted ChIP-qPCR of AR at the *KLK3*, *KLK2*, *TMPRSS2*, and *FKBP5* regulatory regions in LNCaP (A) and DuCaP (B) cells expressing an empty vector (Vec.), wild-type METTL3 (M3-WT), catalytically inactive METTL3 (M3-CD), wild-type NUP93 (N93-WT), or the METTL3-binding-defective NUP93 mutant (N93-AP) under hormone-depleted conditions. The *KIAA0066* locus was included as a negative-control region. AR enrichment is presented as the percentage of input chromatin. Data are presented as mean ± SD from three independent experiments. *P* values were determined by two-tailed paired Student’s *t* tests for the indicated comparisons.

**Supplemental Figure S7.**
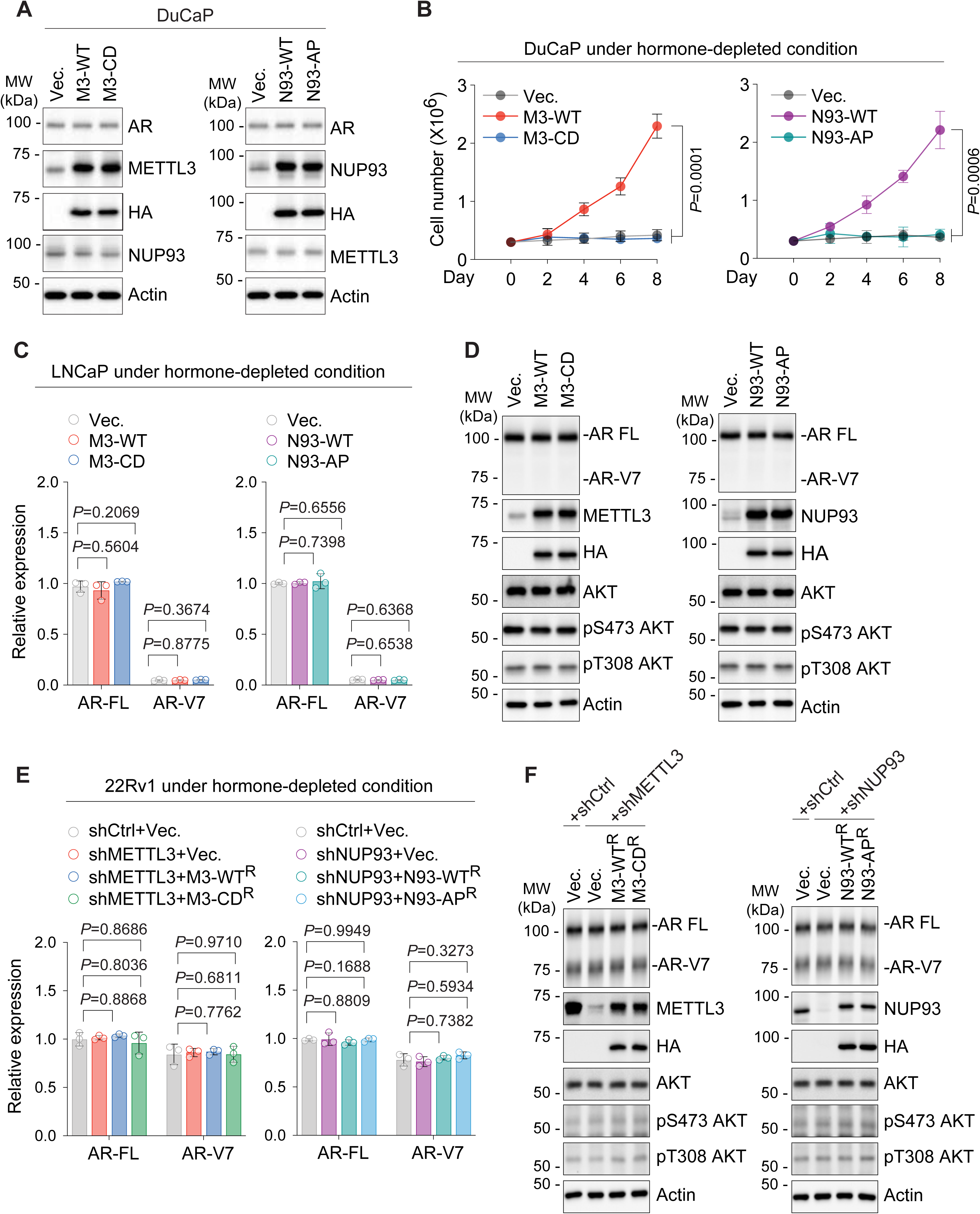
METTL3 and NUP93 promote hormone-independent prostate cancer cell growth without inducing AR-V7 expression or AKT signaling. **(A)** Immunoblot analysis of AR, HA-tagged METTL3 or NUP93, and endogenous NUP93 or METTL3 in DuCaP cells expressing an empty vector (Vec.), wild-type METTL3 (M3-WT), catalytically inactive METTL3 (M3-CD), wild-type NUP93 (N93-WT), or the METTL3-binding-defective NUP93 mutant (N93-AP). Actin was served as the loading control. Blots are representative of three independent experiments. **(B)** Proliferation of DuCaP cells expressing an empty vector (Vec.), M3-WT, M3-CD, N93-WT, or N93-AP under hormone-depleted conditions. **(C)** RT-qPCR analysis of full-length AR (AR-FL) and AR-V7 expression in LNCaP cells expressing an empty vector (Vec.), M3-WT, M3-CD, N93-WT, or N93-AP under hormone-depleted conditions. Expression levels were normalized to *GAPDH* and are presented relative to those in the corresponding vector-control cells. **(D)** Immunoblot analysis of AR-FL, AR-V7, total AKT, AKT phosphorylated at Ser473 [p-AKT (Ser473)], and AKT phosphorylated at Thr308 [p-AKT (Thr308)] in LNCaP cells expressing an empty vector (Vec.), M3-WT, M3-CD, N93-WT, or N93-AP under hormone-depleted conditions. Expression of the indicated METTL3 and NUP93 constructs was confirmed by immunoblotting for METTL3, NUP93, and HA. Actin was served as the loading control. Blots are representative of three independent experiments. **(E)** RT-qPCR analysis of AR-FL and AR-V7 expression in 22Rv1 cells following endogenous METTL3 or NUP93 depletion and re-expression of the corresponding shRNA-resistant wild-type or mutant proteins under hormone-depleted conditions. Expression levels were normalized to *GAPDH* and are presented relative to those in the corresponding shCtrl cells expressing an empty vector (Vec.). **(F)** Immunoblot analysis of AR-FL, AR-V7, total AKT, p-AKT (Ser473), and p-AKT (Thr308) under the conditions described in (E). METTL3 or NUP93 depletion and re-expression of the corresponding HA-tagged rescue constructs were confirmed by immunoblotting. Actin was served as the loading control. Blots are representative of three independent experiments. Data in (B, C, and E) are presented as mean ± SD from three independent experiments. *P* values were determined by two-tailed paired Student’s *t* tests for the indicated comparisons.

**Supplemental Figure S8.**
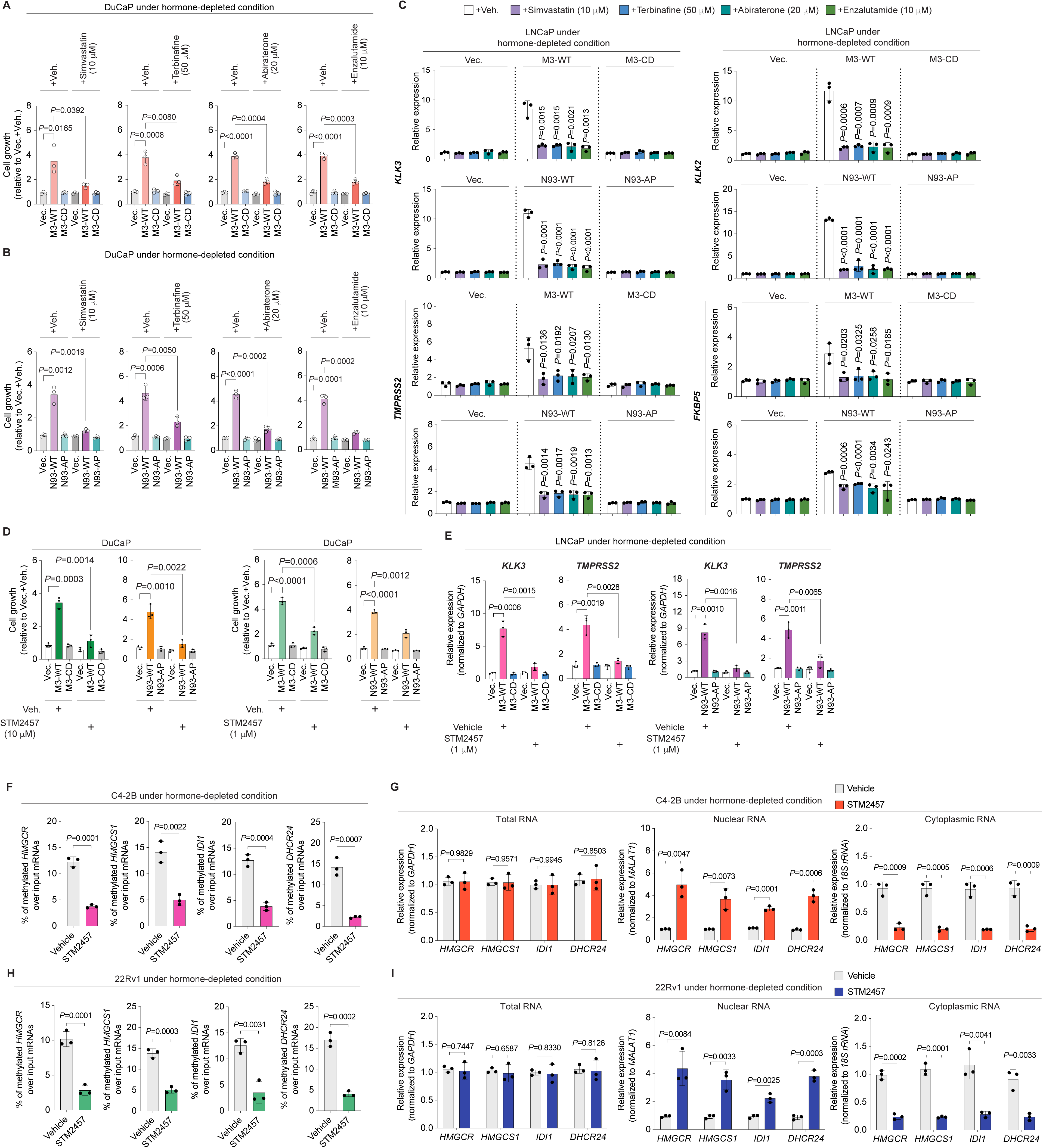
Pharmacologic inhibition of cholesterol biosynthesis, AR signaling, or METTL3 suppresses the malignant phenotypes and mRNA export driven by the m^6^A-METTL3-NUP93 axis. **(A-B)** Cell growth of DuCaP cells expressing an empty vector (Vec.), wild-type METTL3 (M3-WT), or catalytically inactive METTL3 (M3-CD) (A) and an empty vector, wild-type NUP93 (N93-WT), or the METTL3-binding-defective NUP93 mutant (N93-AP) (B) under hormone-depleted conditions. Cells were treated with vehicle, simvastatin (10 μM), terbinafine (50 μM), abiraterone (20 μM), or enzalutamide (10 μM). Cell growth was normalized to that of the corresponding vehicle-treated vector control. **(C)** RT-qPCR analysis of the AR target genes *KLK3*, *KLK2*, *TMPRSS2*, and *FKBP5* in LNCaP cells expressing an empty vector (Vec.), M3-WT, M3-CD, N93-WT, or N93-AP under hormone-depleted conditions. Cells were treated with vehicle, simvastatin (10 μM), terbinafine (50 μM), abiraterone (20 μM), or enzalutamide (10 μM). Gene expression was normalized to *GAPDH* and is presented relative to that in the corresponding vehicle-treated vector-control cells. **(D)** Cell growth of DuCaP cells expressing an empty vector (Vec.), M3-WT, M3-CD, N93-WT, or N93-AP under hormone-depleted conditions following treatment with vehicle or STM2457 at 10 μM (left) or 1 μM (right). Cell growth was normalized to that of the corresponding vehicle-treated vector control. **(E)** RT-qPCR analysis of *KLK3* and *TMPRSS2* expression in LNCaP cells expressing an empty vector (Vec.), M3-WT, M3-CD, N93-WT, or N93-AP under hormone-depleted conditions following treatment with vehicle or STM2457 (1 μM). Gene expression was normalized to *GAPDH* and is presented relative to that in the corresponding vehicle-treated vector-control cells. **(F, H)** m^6^A enrichment on *HMGCR*, *HMGCS1*, *IDI1*, and *DHCR24* mRNAs in C4-2B (F) and 22Rv1 (H) cells treated with vehicle or STM2457 under hormone-depleted conditions. m^6^A enrichment was measured by m^6^A-RIP–qPCR and is presented as the percentage of methylated RNA relative to input. **(G, I)** Total (left), nuclear (middle), and cytoplasmic (right) levels of *HMGCR*, *HMGCS1*, *IDI1*, and *DHCR24* mRNAs in C4-2B (G) and 22Rv1 (I) cells treated with vehicle or STM2457 under hormone-depleted conditions. Total RNA was normalized to *GAPDH*, whereas nuclear and cytoplasmic RNA levels were normalized to *MALAT1* and 18S rRNA, respectively. Data are presented as mean ± SD from three independent experiments. *P* values were determined by two-tailed paired Student’s *t* tests for the indicated comparisons.

## Methods

### Cell Lines and Culture Conditions

Human prostate cancer cell lines LNCaP, DuCaP, LNCaP-abl, C4-2B, and 22Rv1 and the benign prostate epithelial line RWPE-1 and LHSAR were used in this study. LNCaP, DuCaP, C4-2B, and 22Rv1 cells were maintained in RPMI 1640 supplemented with 10% fetal bovine serum (FBS) and 1% penicillin-streptomycin (P/S). LNCaP-abl cells were maintained in phenol-red free RPMI 1640 supplemented with 10% charcoal-stripped fetal bovine serum and 1% penicillin-streptomycin. RWPE-1 cells were maintained in keratinocyte Serum-Free Medium (K-SFM) supplemented with bovine pituitary extract and recombinant epidermal growth factor according to the supplier’s instructions. LHSAR cells were maintained in PrEBM Basal Medium with PrEGMTM SingleQuots® Supplements according to the supplier’s instructions. HEK293T cells used for lentiviral production were grown in Dulbecco’s modified Eagle’s medium (DMEM) containing 10% fetal bovine serum and 1% penicillin-streptomycin. Cultures were maintained at 37°C in a humidified atmosphere containing 5% CO_2_. Hormone-depleted experiments were performed after cells were adapted to phenol-red-free medium supplemented with charcoal-stripped serum at least for 4 days. Where indicated, cells received dihydrotestosterone (DHT), simvastatin, terbinafine, abiraterone, enzalutamide, or STM2457 at the indicated concentrations. Vehicle-treated cultures received the corresponding solvent at the same final concentration. All cell lines were routinely tested for mycoplasma contamination using the Mycoplasma Detection Kit–DigitalTest v2.0 (BioTool, catalog no. B39132). These cell lines were not listed in the database of commonly misidentified cell lines maintained by the International Cell Line Authentication Committee.

### Cell Proliferation Assay

Cells were seeded in 60-mm culture dishes at an initial density of 4 x 10^5^ to 6 x 10^5^ cells per dish. Viable cells were counted on the indicated days using a hemocytometer or a Cytation 5 multimode imaging reader (BioTek). For drug-treatment experiments, cells were treated with the indicated compounds at the concentrations and durations specified in the corresponding figure legends.

### Human Tissue Microarrays and Immunohistochemistry

Prostate tissue microarrays (TMAs) containing benign prostate, primary prostate adenocarcinoma, and castration-resistant prostate cancer specimens were obtained from Dr. Colm Morrissey at the University of Washington. TMA sections (5 μm thick) were deparaffinized in xylene and rehydrated through a graded ethanol series. Heat-induced antigen retrieval was performed in 10 mmol/L citrate buffer (pH 6.0). Endogenous peroxidase activity and endogenous avidin and biotin were blocked using reagents from Vector Laboratories. Sections were subsequently blocked with 5% normal goat–horse–chicken serum overnight at 4°C. TMA sections were incubated with primary antibodies against METTL3 (Proteintech, 15073-1-AP; 1:500) or NUP93 (Atlas antibodies, HOA017937; 1:500), followed by the appropriate biotinylated secondary antibody and avidin–biotin complex reagent (Vector Laboratories). Immunoreactivity was visualized using Stable DAB (Thermo Fisher Scientific) according to the manufacturer’s instructions. Sections were counterstained with hematoxylin and mounted with Cytoseal XYL (Richard-Allan Scientific). Rabbit IgG was used as a negative control.

### Quantification of Immunohistochemical Staining

METTL3 immunoreactivity was evaluated by a pathologist who was blinded to the sample category and clinical information. IHC staining was assessed using a combination of both intensity (score 0, no staining; score 1, weak; score 2, moderate; and score 3, strong) and proportion of stained cells (score 0, 0 to 50%; score 1, 51 to 60%; score 2, 61 to 70%; score 3, 71 to 80%; score 4, 81 to 90%; and score 5, 91 to 100%). The final score was obtained by multiplying the intensity and the proportion scores, resulting in a value ranging from 0 to 15. Cores that were missing, damaged, or contained insufficient evaluable tissue were excluded from the analysis.

Specimens were classified as having high or low protein levels based on percentile-ranked staining scores, with the top 65% classified as high expression and the remaining 35% as low expression. For the joint expression level analysis, only cases with staining data available for both proteins were included and were assigned to one of the following four expression categories: METTL3-high/NUP93-high, METTL3-high/NUP93-low, METTL3-low/NUP93-high, or METTL3-low/NUP93-low. Differences in the frequency of each expression category between castration-resistant prostate cancer and normal adjacent tissues or primary prostate cancer were evaluated using two-sided Fisher’s exact tests, with *P* < 0.05 considered statistically significant.

### Plasmid Construction and sgRNA Construction

Lentiviral constructs expressing wild-type METTL3 (M3-WT), catalytically inactive METTL3 (M3-CD), wild-type NUP93 (N93-WT), and the METTL3-binding-defective NUP93 A333P mutant (N93-AP), including the corresponding shRNA-resistant rescue constructs, were described previously(23). The dCas13-based RNA m^6^A-editing constructs, including nuclear-localized ALKBH5 and METTL3 fusion proteins and their corresponding catalytically inactive derivatives, were also described previously(23). For the present study, sgRNAs targeting m^6^A-enriched regions of the *HMGCR* and *DHCR24* transcripts were designed on the basis of the m^6^A-seq profiles generated in prostate cancer cells. The corresponding oligonucleotides were synthesized, annealed, and cloned into the Cas13 sgRNA expression vector (Addgene, #103854). A non-targeting sgRNA was used as the negative control. All sgRNA constructs were verified by Sanger sequencing. Oligonucleotide sequences for sgRNA construction are provided in **Supplemental Table S1**.

### Lentivirus Production and Generation of Stable Cell Lines

Lentiviral particles were produced by cotransfecting HEK293T cells with the indicated lentiviral transfer vector, a packaging plasmid (pCMV-psPAX2), and the VSV-G envelope plasmid. Virus-containing medium was collected at 48 and 72 hours after transfection, passed through a 0.45-μm filter, and concentrated using Lenti-X Concentrator (Takara Bio). Target cells were transduced with the concentrated viral particles in the presence of polybrene (5 μg/mL; Sigma-Aldrich). At 24 hours after transduction, cells were selected with puromycin (1 μg/mL; Sigma-Aldrich) or hygromycin B (100–200 μg/mL; Sigma-Aldrich), as appropriate for the vector. Selection was maintained for two to three passages before the cells were used in subsequent experiments. For depletion-rescue experiments, cells were sequentially transduced with lentiviruses expressing an shRNA-resistant wild-type or mutant rescue construct and the corresponding shRNA targeting endogenous METTL3 or NUP93. Depletion of the endogenous protein and expression of the rescue protein were confirmed by immunoblotting before functional analysis.

### RNA Interference and Transient Transfection

Cells were transfected with control or gene-targeting siRNAs using Lipofectamine RNAiMAX Transfection Reagent (Thermo Fisher Scientific) according to the manufacturer’s instructions. Plasmids and sgRNA expression vectors were transfected using Lipofectamine 3000 (Thermo Fisher Scientific). For dCas13-based RNA editing, cells were cotransfected with the indicated sgRNA and dCas13 fusion-protein expression plasmid at a mass ratio of 1:3. Cells were collected 72 hours after transfection unless otherwise indicated. Knockdown or RNA-editing efficiency was confirmed in the same experimental samples used for downstream analyses.

### RNA Isolation, Nuclear-Cytoplasmic Fractionation, and RT-qPCR

RNA isolation and reverse transcription–quantitative PCR (RT-qPCR) were performed as previously described(23,63). For total RNA analysis, RNA was extracted using TRIzol Reagent (Thermo Fisher Scientific). For nuclear-cytoplasmic fractionation, cells at approximately 80% confluence were washed twice with ice-cold PBS and fractionated using NE-PER Nuclear and Cytoplasmic Extraction Reagents (Thermo Fisher Scientific) according to the manufacturer’s instructions. RNA from the nuclear fraction was extracted using TRIzol LS Reagent, whereas cytoplasmic RNA was isolated using TRIzol Reagent. Fractionated RNA was treated with TURBO DNase (Thermo Fisher Scientific). RNA concentration was measured using a Cytation 5 multimode reader (BioTek), and 1 μg of RNA was reverse-transcribed using the High-Capacity cDNA Reverse Transcription Kit (Thermo Fisher Scientific). Quantitative PCR was performed using SYBR Green qPCR Master Mix (Bio-Rad). Relative RNA abundance was calculated using the comparative Ct method. Total RNA was normalized to *GAPDH*, whereas nuclear and cytoplasmic RNA were normalized to *MALAT1* and *18S rRNA*, respectively. Primer sequences for RT-qPCR are provided in **Supplemental Table S2**.

### m^6^A RNA Immunoprecipitation and qPCR (m^6^A MeRIP-qPCR)

RNA was chemically fragmented in 100 mmol/L Tris-HCl (pH 7.4) and 100 mmol/L ZnCl_2_ at 70 degrees C for 15 minutes. Fragmented RNA was incubated for 4 hours at 4°C with anti-m^6^A antibody (Millipore, ABE572) prebound to a 1:1 mixture of Protein A and Protein G magnetic beads in immunoprecipitation buffer containing 150 mmol/L NaCl, 10 mmol/L Tris-HCl (pH 7.5), 0.1% IGEPAL CA-630, and RNase inhibitor. Complexes were washed twice each with the original buffer, a 50 mmol/L NaCl buffer, and a 500 mmol/L NaCl buffer. Bound RNA was released in RLT buffer and purified on an RNeasy MinElute column. For gene-specific assays, 10% of fragmented RNA was retained as input. Enrichment at HMGCR, HMGCS1, IDI1, and DHCR24 regions was quantified by RT-qPCR and expressed relative to input. Primer sequences for MeRIP-qPCR are provided in **Supplemental Table S2**.

### m^6^A-seq and Data Analysis

m^6^A-seq libraries were generated from immunoprecipitated RNA and matched input using the SMARTer Stranded Total RNA-Seq Kit v2-Pico Input Mammalian and sequenced on an Illumina platform with paired-end reads. Two independent biological replicates were analyzed per condition. Adapters and low-quality terminal bases were removed with Cutadapt, and reads of at least 25 nucleotides were aligned to the human reference genome hg19 with STAR. PCR duplicates were removed with Picard. Coverage tracks were normalized to 10 million mapped reads. m^6^A-enriched regions were identified from immunoprecipitated and input libraries using MeTDiff. Peaks were assigned to annotated transcripts, and genome-browser tracks were displayed in Integrative Genomics Viewer.

### Fractionation RNA-seq and Data Analysis

Total and nuclear RNA libraries were prepared with the KAPA Stranded mRNA-Seq Kit. ERCC spike-in RNA was added to nuclear samples at a 1:100 dilution before library preparation. Reads were aligned to the human reference genome hg19 with STAR. For spike-in-containing samples, the reference included the ERCC sequences. Uniquely mapped reads were counted with featureCounts, and genes with fewer than one read in at least two libraries were removed. Size factors for nuclear RNA were estimated from ERCC features with more than 10 reads in over half of the libraries. Differential abundance was tested with DESeq2. Transcripts were considered significantly changed at an absolute fold change of at least 2 and *P* <1 x 10^-10^. Nuclear-retained candidates were defined as transcripts increased in the nuclear fraction without a corresponding change in total RNA and were intersected with transcripts carrying an m^6^A peak. The overlap between METTL3- and NUP93-dependent gene sets was evaluated with a hypergeometric test. Pathway enrichment was performed against the MSigDB Hallmark collection using GSEA. Multiple-testing correction used the Benjamini-Hochberg procedure. Gene sets meeting an adjusted *P* value or false-discovery rate (FDR) < 0.01 were considered enriched.

### Immunoblotting and Co-immunoprecipitation

For immunoblotting, cells were collected and lysed in buffer containing 0.5% IGEPAL CA-630, 0.5% Triton X-100, 150 mmol/L NaCl, 2 mmol/L EDTA (pH 8.0), 50 mmol/L Tris-HCl (pH 7.5), 10 mmol/L sodium fluoride, 2 mmol/L sodium orthovanadate, 1 mmol/L phenylmethylsulfonyl fluoride, and 1× protease inhibitor cocktail (Roche). Protein concentrations were determined using the Pierce BCA Protein Assay Kit (Thermo Fisher Scientific). Equal amounts of protein were separated by SDS-PAGE and transferred to PVDF membranes. Membranes were blocked with 5% nonfat milk in Tris-buffered saline containing 0.02% Tween 20 (TBS-T) for 1 hour at room temperature and incubated with the indicated primary antibodies overnight at 4°C. After washing, membranes were incubated with the appropriate horseradish peroxidase–conjugated secondary antibodies and developed using enhanced chemiluminescence. For co-immunoprecipitation, cell lysates were precleared with Protein A/G Plus Agarose (Santa Cruz Biotechnology) and then incubated overnight at 4°C with the indicated primary antibody and Dynabeads Protein A or Protein G magnetic beads (Thermo Fisher Scientific). Immunoprecipitates were washed three times with lysis buffer, resolved by SDS-PAGE, and analyzed by immunoblotting. Antibody information is provided in **Supplemental Table S3**.

### Proximity Ligation Assay

Proximity ligation assays were performed using the Duolink In Situ Red Starter Kit Mouse/Rabbit (Sigma-Aldrich, DUO92101) according to the manufacturer’s instructions. Cells grown on glass coverslips were fixed with 3.7% formaldehyde in PBS for 10 minutes at room temperature and permeabilized with 0.3% Triton X-100 in PBS for 10 minutes. Cells were blocked with Duolink Blocking Solution for 60 minutes at 37°C in a humidified chamber and then incubated overnight at 4°C with a rabbit polyclonal antibody against METTL3 (Proteintech, 15073-1-AP) and a mouse monoclonal antibody against NUP93 (E-8; Santa Cruz Biotechnology, sc-374399), diluted in Duolink Antibody Diluent. After two 5-minute washes with Duolink Wash Buffer A, cells were incubated with anti-rabbit PLUS and anti-mouse MINUS proximity probes for 100 minutes at 37°C in a humidified chamber. Cells were washed twice with Wash Buffer A and incubated with ligase in ligation buffer for 30 minutes at 37°C. Following two additional washes with Wash Buffer A, rolling-circle amplification was performed using the supplied polymerase and amplification buffer containing fluorescent detection probes according to the manufacturer’s instructions. Coverslips were subsequently washed with Duolink Wash Buffer B and mounted using Duolink In Situ Mounting Medium containing DAPI. Images were acquired using a ZEISS LSM 980 confocal microscope equipped with Airyscan 2 under identical acquisition settings for all experimental groups. Airyscan image processing was performed using ZEN software (ZEISS), with the same processing parameters applied across all groups.

### Soft-Agar Colony-Formation Assay

Viable cells were suspended in 0.3% agarose (Sigma-Aldrich, A0701) and plated at a low density in six-well plates containing a base layer of 0.6% agar (Sigma-Aldrich, A1296). LNCaP cells were seeded at 5,000 cells per well. C4-2B and 22Rv1 cells were seeded at 3,000-5,000 cells per well. Cells were maintained under complete or hormone-depleted culture conditions, as indicated. After 18–24 days, colonies were stained with crystal violet (Sigma-Aldrich, C0775), imaged, and quantified using ImageJ.

### Transwell Migration and Invasion Assays

Cell migration and invasion were evaluated using Falcon permeable supports fitted with transparent 8.0-μm-pore PET membranes (Corning, 353097), according to the manufacturer’s instructions. For invasion assays, membranes were coated with Corning Matrigel Basement Membrane Matrix (Corning, 354234); uncoated membranes were used for migration assays. Inserts were placed in companion plates containing complete culture medium supplemented with 10% FBS as a chemoattractant. LNCaP, C4-2B, and 22Rv1 cells were suspended in serum-free medium and seeded into the upper chambers at 5 × 10^4^ cells per insert. After 16–18 hours, cells remaining on the upper surface of the membrane were removed. Cells that had migrated or invaded through the membrane were fixed with 3.7% formaldehyde, permeabilized with 100% methanol, and stained with crystal violet (Sigma-Aldrich, C0775). Representative images were acquired using a Cytation 5 multimode reader (BioTek), and migrated or invaded cells were quantified using ImageJ.

### Cellular Cholesterol Measurement

Total cellular cholesterol was measured using the Cholesterol/Cholesteryl Ester Quantitation Assay Kit (Abcam, ab65359) according to the manufacturer’s instructions. Approximately 1 × 10^6^ cells were collected, washed with ice-cold PBS, and resuspended in 200 μL of chloroform:isopropanol:NP-40 (7:11:0.1) to extract cellular lipids. After centrifugation at 15,000 × *g* for 5–10 minutes, the organic phase was transferred to a new tube, air-dried at 50°C, and placed under vacuum for 30 minutes to remove residual solvent. Dried lipids were resuspended in 200 μL of Cholesterol Assay Buffer. Samples and cholesterol standards were incubated with the supplied enzyme mix, cholesterol esterase, and probe for 60 minutes at 37°C, protected from light. Absorbance was measured at 570 nm using a Cytation 5 multimode reader (BioTek). Total cholesterol concentrations were calculated from the standard curve and normalized to the corresponding control group.

### Cholesterol-Uptake Assay

Cellular cholesterol uptake was measured using the Cholesterol Uptake Assay Kit (Abcam, ab236212) according to the manufacturer’s instructions. Cells were incubated for 24 hours in serum-free medium containing 20 μg/mL NBD cholesterol. The NBD cholesterol–containing medium was then removed, and cells were washed with the supplied assay buffer to remove extracellular fluorescence. Cellular fluorescence was measured using a Cytation 5 multimode reader (BioTek) at excitation and emission wavelengths of 485 and 535 nm, respectively. Fluorescence values were corrected for background and normalized to the corresponding control group.

### Dihydrotestosterone Measurement

Intracellular dihydrotestosterone (DHT) levels were measured using a competitive DHT ELISA kit (MyBioSource, MBS2700128) according to the manufacturer’s instructions. Adherent cells were washed with ice-cold PBS, detached, and collected by centrifugation at 1,000 × *g* for 5 minutes. Cell pellets were washed three times with ice-cold PBS and resuspended in lysis buffer at a density of approximately 1 × 10^7^ cells/mL. Lysates were clarified by centrifugation at 1,500 × *g* for 10 minutes at 4°C, and the resulting supernatants were used for the assay. Standards and cell lysates were added to the antibody-coated microplate, followed by Detection Reagent A, and incubated for 1 hour at 37°C. After washing, Detection Reagent B was added, and the plate was incubated for 30 minutes at 37°C. Wells were washed five times and incubated with TMB substrate for 10–20 minutes at 37°C while protected from light. The reaction was terminated with Stop Solution, and absorbance was measured immediately at 450 nm using a Cytation 5 multimode reader (BioTek). DHT concentrations were calculated from the standard curve after averaging duplicate measurements and subtracting the blank absorbance. Values were multiplied by the corresponding dilution factor, when applicable, and are reported as pg/mL.

### Luciferase Reporter Assay

Androgen receptor transcriptional activity was evaluated using a firefly luciferase reporter containing KLK3 enhancer elements. Cells were cotransfected with the firefly luciferase reporter and a Renilla luciferase plasmid as an internal transfection control. Following transfection, cells were maintained under hormone-depleted conditions and treated as indicated. Firefly and Renilla luciferase activities were measured using the Dual-Luciferase Reporter Assay System (Promega) according to the manufacturer’s instructions. Firefly luciferase activity was normalized to the corresponding Renilla luciferase activity, and the resulting values were expressed relative to the matched vector-control group.

### Chromatin Immunoprecipitation (ChIP)-qPCR

Chromatin immunoprecipitation followed by quantitative PCR (ChIP-qPCR) was performed as previously described, with minor modifications(63). Cells were sequentially cross-linked with 6 mmol/L disuccinimidyl glutarate (CovaChem) for 30 minutes and 1% formaldehyde (Sigma-Aldrich) for 10 minutes at room temperature. Nuclei were isolated by sequential extraction with LB1 buffer containing 50 mmol/L HEPES-KOH (pH 7.5), 140 mmol/L NaCl, 1 mmol/L EDTA (pH 8.0), 10% glycerol, 0.5% NP-40, 0.25% Triton X-100, and 1× cOmplete Protease Inhibitor Cocktail; LB2 buffer containing 10 mmol/L Tris-HCl (pH 8.0), 200 mmol/L NaCl, 1 mmol/L EDTA (pH 8.0), 0.5 mmol/L EGTA (pH 8.0), and 1× cOmplete Protease Inhibitor Cocktail; and LB3 buffer containing 10 mmol/L Tris-HCl (pH 8.0), 100 mmol/L NaCl, 1 mmol/L EDTA (pH 8.0), 0.5 mmol/L EGTA (pH 8.0), 0.1% sodium deoxycholate, 0.5% *N*-lauroylsarcosine, and 1× cOmplete Protease Inhibitor Cocktail. Chromatin was sheared using a Q800R sonicator (Qsonica) and incubated overnight at 4°C with the indicated antibody or corresponding control IgG conjugated to a mixture of Dynabeads Protein A and Protein G magnetic beads (Thermo Fisher Scientific). Immunoprecipitates were washed three times with RIPA wash buffer containing 50 mmol/L HEPES-KOH (pH 7.5), 500 mmol/L LiCl, 1 mmol/L EDTA, 1% NP-40, 0.7% sodium deoxycholate, and 1× cOmplete Protease Inhibitor Cocktail, followed by two washes with TE buffer containing 10 mmol/L Tris-HCl (pH 8.0) and 1 mmol/L EDTA. Chromatin was eluted in buffer containing 1% SDS, 10 mmol/L Tris-HCl (pH 8.0), and 10 mmol/L EDTA. Cross-links were reversed at 65°C overnight, and DNA was treated with RNase A and proteinase K before purification. Enrichment at the indicated genomic regions was measured by quantitative PCR and calculated relative to input DNA. Primer sequences are provided in **Supplemental Table S2**.

### Chromatin Immunoprecipitation Sequencing (ChIP-seq)

AR ChIP-seq was performed using the cross-linking, chromatin preparation, immunoprecipitation, washing, elution, and DNA-purification procedures described above for ChIP-qPCR. Sheared chromatin was immunoprecipitated with an androgen receptor antibody (D6F11; Cell Signaling Technology, #5153). Sequencing libraries were prepared from AR-immunoprecipitated and matched input DNA using the KAPA HyperPrep Kit (Roche) according to the manufacturer’s instructions and sequenced on an Illumina NextSeq platform. Sequencing reads were aligned to the human reference genome (hg19) using Bowtie. AR-binding peaks were called against matched input DNA using MACS with a q-value cutoff of 1 x 10^-5^. Peaks overlapping ENCODE/UCSC blacklist regions were excluded from subsequent analyses. ChIP-seq signals were normalized to 10 million mapped reads per sample. Heatmaps and aggregate profiles were generated across the identified AR-binding sites within ±2 kb of peak centers. Enriched transcription-factor binding motifs were identified using HOMER within ±300 bp of peak summits. Genome-browser tracks were visualized using the Integrative Genomics Viewer (IGV).

### In vivo tumor growth in xenograft mouse model of prostate cancer

Five 6-week-old male BALB/c nude mice (Charles River Laboratories) were surgically castrated. After complete postoperative recovery, viable 22Rv1 cells (2 × 10^6^) were suspended in serum-free medium containing 50% Corning Matrigel Matrix High Concentration (Corning, 08-774-391) and injected subcutaneously into the lower flanks. To control for differences in the baseline proliferative capacity of the engineered cell populations, paired cell populations were implanted into opposite flanks of the same mouse. Control cells expressing control shRNA and empty vector (Vec. + shCtrl) and NUP93-depleted cells expressing an empty vector (shNUP93 + Vec.) were implanted into the left and right flanks, respectively. NUP93-depleted cells reconstituted with shRNA-resistant wild-type NUP93 (shNUP93 + N93-WT) and those reconstituted with the shRNA-resistant METTL3-binding-defective NUP93 A333P mutant (shNUP93 + N93-AP) were implanted into the left and right flanks, respectively. Tumor dimensions and body weights were measured every 3 days throughout the study. Tumor volume was calculated using the formula length (L) x width (W)^2^/2, where L and W represent the longest and shortest tumor diameters, respectively. Mice were monitored for changes in body weight and general condition as indicators of treatment-associated toxicity.

### Patient-Derived Xenograft Study

LuCaP 235.2 castration-resistant prostate cancer patient-derived xenografts were propagated subcutaneously in castrated male C.B-17 SCID mice (Charles River Laboratories). Similarly sized tumor fragments were implanted subcutaneously using a trocar. When tumors reached approximately 200 mm^3^, mice were randomly assigned to receive vehicle or STM2457 (50 mg/kg) by intraperitoneal (IP) injection. Treatment was administered for 5 consecutive days followed by a 2-day treatment-free interval, and this schedule was repeated throughout the study. Tumor dimensions and body weights were measured every 3 days. Tumor volume was calculated as L x W^2^/2 as described above.

### Statistical Analysis

Statistical analyses were performed using GraphPad Prism (GraphPad Software). Unless otherwise indicated, quantitative data are presented as mean ± SD from three independent biological experiments. Tumor growth and mouse body-weight data are presented as mean ± SEM. Comparisons between two groups were performed using two-tailed Student’s *t* tests. Paired t tests were used for experimentally matched samples, including matched human tissues and paired xenografts implanted into opposite flanks of the same mouse. Unpaired t tests were used for comparisons between independent groups, including vehicle- and STM2457-treated patient-derived xenografts. The statistical test and definition of n are provided in the corresponding figure legends. Exact *P* values are reported in the figures, and *P* < 0.05 was considered statistically significant. For sequencing-based analyses, multiple-hypothesis testing was controlled using the Benjamini–Hochberg procedure where indicated. The statistical methods and significance thresholds applied to RNA-seq, m^6^A-seq, ChIP-seq, pathway-enrichment, and gene-set overlap analyses are described in the corresponding Methods sections. No statistical method was used to predetermine sample size, and no data points were excluded unless otherwise indicated.

Data and materials availability: All the genome-wide datasets generated in this study, including RNA-seq and ChIP-seq, have been deposited at the Gene Expression Omnibus database (http://ncbi.nlm.nih.gov/geo/) with an accession number GSE345532 and GSE345531, respectively. Previously published nuclear RNA-seq, total RNA-seq, and m^6^A-seq datasets used in this study are available under accession number GSE248589. The cohort studies of patients with prostate cancer are all from Oncomine with GEO accession numbers available as GSE35598, GSE55945, and GSE6919. The CRISPR-Cas9 knockout screening data are retrieved from DepMap.

## Acknowledgments

This work was supported by the National Cancer Institute of the National Institutes of Health under award numbers R01CA279696 (to J.L.) and R01CA279681 (to K.X.). The authors thank the Optical Imaging Facility at UT Health San Antonio for assistance with confocal microscopy. The ZEISS LSM 980 confocal microscope with Airyscan 2 was funded by NIH S10 grant 1S10OD036297-01. The Optical Imaging Facility is supported by UT Health San Antonio and the NIH/NCI Cancer Center Support Grant P30 CA054174. The establishment and characterization of the LuCaP PDX models was supported by the Pacific Northwest Prostate Cancer SPORE (P50CA097186), the P01 NIH grant (P01CA163227) and the Institute of Prostate Cancer Research.

## Conflict of Interest

E.C. served as a paid consultant to DotQuant, and received Institutional sponsored research funding unrelated to this work from Astra Zeneca, AbbVie, Gilead, Sanofi, Zenith Epigenetics, Bayer Pharmaceuticals, Forma Therapeutics, Genentech, GSK, Janssen Research, Kronos Bio, Foghorn Therapeutics, K36 Therapeutics, BoundlessBio, and MacroGenics.

## References

1. Simon DN, Rout MP. Cancer and the nuclear pore complex. Adv Exp Med Biol 2014;773:285–307 doi 10.1007/978-1-4899-8032-8_13.

2. Sakuma S, D’Angelo MA. The roles of the nuclear pore complex in cellular dysfunction, aging and disease. Semin Cell Dev Biol 2017;68:72–84 doi 10.1016/j.semcdb.2017.05.006.

3. Kuusisto HV, Wagstaff KM, Alvisi G, Roth DM, Jans DA. Global enhancement of nuclear localization-dependent nuclear transport in transformed cells. FASEB J 2012;26(3):1181–93 doi 10.1096/fj.11-191585.

4. Lewin JM, Lwaleed BA, Cooper AJ, Birch BR. The direct effect of nuclear pores on nuclear chemotherapeutic concentration in multidrug resistant bladder cancer: the nuclear sparing phenomenon. J Urol 2007;177(4):1526–30 doi 10.1016/j.juro.2006.11.048.

5. Kabachinski G, Schwartz TU. The nuclear pore complex--structure and function at a glance. J Cell Sci 2015;128(3):423–9 doi 10.1242/jcs.083246.

6. Hegedusova E, Marsalova V, Kulkarni S, Paris Z. Trafficking and/or division: Distinct roles of nucleoporins based on their location within the nuclear pore complex. RNA Biol 2022;19(1):650–61 doi 10.1080/15476286.2022.2067711.

7. Nakamura T, Largaespada DA, Lee MP, Johnson LA, Ohyashiki K, Toyama K, et al. Fusion of the nucleoporin gene NUP98 to HOXA9 by the chromosome translocation t(7;11)(p15;p15) in human myeloid leukaemia. Nat Genet 1996;12(2):154–8 doi 10.1038/ng0296-154.

8. Van Vlierberghe P, van Grotel M, Tchinda J, Lee C, Beverloo HB, van der Spek PJ, et al. The recurrent SET-NUP214 fusion as a new HOXA activation mechanism in pediatric T-cell acute lymphoblastic leukemia. Blood 2008;111(9):4668–80 doi 10.1182/blood-2007-09-111872.

9. Alanee S, Delfino K, Wilber A, Robinson K, Brard L, Semaan A. Single nucleotide variant in Nucleoporin 107 may be predictive of sensitivity to chemotherapy in patients with ovarian cancer. Pharmacogenet Genomics 2017;27(7):264–9 doi 10.1097/FPC.0000000000000288.

10. Nataraj NB, Noronha A, Lee JS, Ghosh S, Mohan Raju HR, Sekar A, et al. Nucleoporin-93 reveals a common feature of aggressive breast cancers: robust nucleocytoplasmic transport of transcription factors. Cell Rep 2022;38(8):110418 doi 10.1016/j.celrep.2022.110418.

11. Nofrini V, Di Giacomo D, Mecucci C. Nucleoporin genes in human diseases. Eur J Hum Genet 2016;24(10):1388–95 doi 10.1038/ejhg.2016.25.

12. Xylourgidis N, Fornerod M. Acting out of character: regulatory roles of nuclear pore complex proteins. Dev Cell 2009;17(5):617–25 doi 10.1016/j.devcel.2009.10.015.

13. Raices M, D’Angelo MA. Nuclear pore complexes and regulation of gene expression. Curr Opin Cell Biol 2017;46:26–32 doi 10.1016/j.ceb.2016.12.006.

14. Lesbirel S, Wilson SA. The m(6)A-methylase complex and mRNA export. Biochim Biophys Acta Gene Regul Mech 2019;1862(3):319–28 doi 10.1016/j.bbagrm.2018.09.008.

15. Roundtree IA, Luo GZ, Zhang Z, Wang X, Zhou T, Cui Y, et al. YTHDC1 mediates nuclear export of N(6)-methyladenosine methylated mRNAs. Elife 2017;6 doi 10.7554/eLife.31311.

16. Edens BM, Vissers C, Su J, Arumugam S, Xu Z, Shi H, et al. FMRP Modulates Neural Differentiation through m(6)A-Dependent mRNA Nuclear Export. Cell Rep 2019;28(4):845–54 e5 doi 10.1016/j.celrep.2019.06.072.

17. Liu J, Yue Y, Han D, Wang X, Fu Y, Zhang L, et al. A METTL3-METTL14 complex mediates mammalian nuclear RNA N6-adenosine methylation. Nat Chem Biol 2014;10(2):93–5 doi 10.1038/nchembio.1432.

18. Ping XL, Sun BF, Wang L, Xiao W, Yang X, Wang WJ, et al. Mammalian WTAP is a regulatory subunit of the RNA N6-methyladenosine methyltransferase. Cell Res 2014;24(2):177–89 doi 10.1038/cr.2014.3.

19. Jia G, Fu Y, Zhao X, Dai Q, Zheng G, Yang Y, et al. N6-methyladenosine in nuclear RNA is a major substrate of the obesity-associated FTO. Nat Chem Biol 2011;7(12):885–7 doi 10.1038/nchembio.687.

20. Zheng G, Dahl JA, Niu Y, Fedorcsak P, Huang CM, Li CJ, et al. ALKBH5 is a mammalian RNA demethylase that impacts RNA metabolism and mouse fertility. Mol Cell 2013;49(1):18–29 doi 10.1016/j.molcel.2012.10.015.

21. Deng X, Qing Y, Horne D, Huang H, Chen J. The roles and implications of RNA m(6)A modification in cancer. Nat Rev Clin Oncol 2023;20(8):507–26 doi 10.1038/s41571-023-00774-x.

22. Lesbirel S, Viphakone N, Parker M, Parker J, Heath C, Sudbery I, et al. The m(6)A-methylase complex recruits TREX and regulates mRNA export. Sci Rep 2018;8(1):13827 doi 10.1038/s41598-018-32310-8.

23. Lee JH, Tingey M, Zhang Z, Buerger F, Hong J, Zhang G, et al. N(6)-adenosine methylation enhances nuclear mRNA export through METTL3 and NUP93. Nat Cell Biol 2026;28(3):553–66 doi 10.1038/s41556-026-01882-3.

24. Rebello RJ, Oing C, Knudsen KE, Loeb S, Johnson DC, Reiter RE, et al. Prostate cancer. Nat Rev Dis Primers 2021;7(1):9 doi 10.1038/s41572-020-00243-0.

25. Leon CG, Locke JA, Adomat HH, Etinger SL, Twiddy AL, Neumann RD, et al. Alterations in cholesterol regulation contribute to the production of intratumoral androgens during progression to castration-resistant prostate cancer in a mouse xenograft model. Prostate 2010;70(4):390–400 doi 10.1002/pros.21072.

26. Mostaghel EA, Solomon KR, Pelton K, Freeman MR, Montgomery RB. Impact of circulating cholesterol levels on growth and intratumoral androgen concentration of prostate tumors. PLoS One 2012;7(1):e30062 doi 10.1371/journal.pone.0030062.

27. Gordon JA, Midha A, Szeitz A, Ghaffari M, Adomat HH, Guo Y, et al. Oral simvastatin administration delays castration-resistant progression and reduces intratumoral steroidogenesis of LNCaP prostate cancer xenografts. Prostate Cancer Prostatic Dis 2016;19(1):21–7 doi 10.1038/pcan.2015.37.

28. Kong Y, Cheng L, Mao F, Zhang Z, Zhang Y, Farah E, et al. Inhibition of cholesterol biosynthesis overcomes enzalutamide resistance in castration-resistant prostate cancer (CRPC). J Biol Chem 2018;293(37):14328–41 doi 10.1074/jbc.RA118.004442.

29. Ashida S, Kawada C, Inoue K. Stromal regulation of prostate cancer cell growth by mevalonate pathway enzymes HMGCS1 and HMGCR. Oncol Lett 2017;14(6):6533–42 doi 10.3892/ol.2017.7025.

30. Han W, Gao S, Barrett D, Ahmed M, Han D, Macoska JA, et al. Reactivation of androgen receptor-regulated lipid biosynthesis drives the progression of castration-resistant prostate cancer. Oncogene 2018;37(6):710–21 doi 10.1038/onc.2017.385.

31. Kohler A, Hurt E. Gene regulation by nucleoporins and links to cancer. Mol Cell 2010;38(1):6–15 doi 10.1016/j.molcel.2010.01.040.

32. Borden KLB. The Nuclear Pore Complex and mRNA Export in Cancer. Cancers (Basel) 2020;13(1) doi 10.3390/cancers13010042.

33. Yang Y, Guo L, Chen L, Gong B, Jia D, Sun Q. Nuclear transport proteins: structure, function, and disease relevance. Signal Transduct Target Ther 2023;8(1):425 doi 10.1038/s41392-023-01649-4.

34. Sakuma S, Raices M, Borlido J, Guglielmi V, Zhu EYS, D’Angelo MA. Inhibition of Nuclear Pore Complex Formation Selectively Induces Cancer Cell Death. Cancer Discov 2021;11(1):176–93 doi 10.1158/2159-8290.CD-20-0581.

35. Rodriguez-Bravo V, Pippa R, Song WM, Carceles-Cordon M, Dominguez-Andres A, Fujiwara N, et al. Nuclear Pores Promote Lethal Prostate Cancer by Increasing POM121-Driven E2F1, MYC, and AR Nuclear Import. Cell 2018;174(5):1200–15 e20 doi 10.1016/j.cell.2018.07.015.

36. Cai C, Chen S, Ng P, Bubley GJ, Nelson PS, Mostaghel EA, et al. Intratumoral de novo steroid synthesis activates androgen receptor in castration-resistant prostate cancer and is upregulated by treatment with CYP17A1 inhibitors. Cancer Res 2011;71(20):6503–13 doi 10.1158/0008-5472.CAN-11-0532.

37. Montgomery RB, Mostaghel EA, Vessella R, Hess DL, Kalhorn TF, Higano CS, et al. Maintenance of intratumoral androgens in metastatic prostate cancer: a mechanism for castration-resistant tumor growth. Cancer Res 2008;68(11):4447–54 doi 10.1158/0008-5472.CAN-08-0249.

38. Dillard PR, Lin MF, Khan SA. Androgen-independent prostate cancer cells acquire the complete steroidogenic potential of synthesizing testosterone from cholesterol. Mol Cell Endocrinol 2008;295(1-2):115–20 doi 10.1016/j.mce.2008.08.013.

39. Hieronymus H, Lamb J, Ross KN, Peng XP, Clement C, Rodina A, et al. Gene expression signature-based chemical genomic prediction identifies a novel class of HSP90 pathway modulators. Cancer Cell 2006;10(4):321–30 doi 10.1016/j.ccr.2006.09.005.

40. Kim JH, Cox ME, Wasan KM. Effect of simvastatin on castration-resistant prostate cancer cells. Lipids Health Dis 2014;13:56 doi 10.1186/1476-511X-13-56.

41. Padyana AK, Gross S, Jin L, Cianchetta G, Narayanaswamy R, Wang F, et al. Structure and inhibition mechanism of the catalytic domain of human squalene epoxidase. Nat Commun 2019;10(1):97 doi 10.1038/s41467-018-07928-x.

42. Attard G, Reid AH, Yap TA, Raynaud F, Dowsett M, Settatree S, et al. Phase I clinical trial of a selective inhibitor of CYP17, abiraterone acetate, confirms that castration-resistant prostate cancer commonly remains hormone driven. J Clin Oncol 2008;26(28):4563–71 doi 10.1200/JCO.2007.15.9749.

43. Tran C, Ouk S, Clegg NJ, Chen Y, Watson PA, Arora V, et al. Development of a second-generation antiandrogen for treatment of advanced prostate cancer. Science 2009;324(5928):787–90 doi 10.1126/science.1168175.

44. Wang S, Lv W, Li T, Zhang S, Wang H, Li X, et al. Dynamic regulation and functions of mRNA m6A modification. Cancer Cell Int 2022;22(1):48 doi 10.1186/s12935-022-02452-x.

45. Miller WL, Auchus RJ. The molecular biology, biochemistry, and physiology of human steroidogenesis and its disorders. Endocr Rev 2011;32(1):81–151 doi 10.1210/er.2010-0013.

46. Fornerod M, Ohno M, Yoshida M, Mattaj IW. CRM1 is an export receptor for leucine-rich nuclear export signals. Cell 1997;90(6):1051–60 doi 10.1016/s0092-8674(00)80371-2.

47. Booth DS, Cheng Y, Frankel AD. The export receptor Crm1 forms a dimer to promote nuclear export of HIV RNA. Elife 2014;3:e04121 doi 10.7554/eLife.04121.

48. Kim WK, Buckley AJ, Lee DH, Hiroto A, Nenninger CH, Olson AW, et al. Androgen deprivation induces double-null prostate cancer via aberrant nuclear export and ribosomal biogenesis through HGF and Wnt activation. Nat Commun 2024;15(1):1231 doi 10.1038/s41467-024-45489-4.

49. Wei XX, Siegel AP, Aggarwal R, Lin AM, Friedlander TW, Fong L, et al. A Phase II Trial of Selinexor, an Oral Selective Inhibitor of Nuclear Export Compound, in Abiraterone-and/or Enzalutamide-Refractory Metastatic Castration-Resistant Prostate Cancer. Oncologist 2018;23(6):656–e64 doi 10.1634/theoncologist.2017-0624.

50. Pelton K, Freeman MR, Solomon KR. Cholesterol and prostate cancer. Curr Opin Pharmacol 2012;12(6):751–9 doi 10.1016/j.coph.2012.07.006.

51. Schaffner CP. Prostatic cholesterol metabolism: regulation and alteration. Prog Clin Biol Res 1981;75A:279–324.

52. Guillaumond F, Bidaut G, Ouaissi M, Servais S, Gouirand V, Olivares O, et al. Cholesterol uptake disruption, in association with chemotherapy, is a promising combined metabolic therapy for pancreatic adenocarcinoma. Proc Natl Acad Sci U S A 2015;112(8):2473–8 doi 10.1073/pnas.1421601112.

53. Raftopulos NL, Washaya TC, Niederprum A, Egert A, Hakeem-Sanni MF, Varney B, et al. Prostate cancer cell proliferation is influenced by LDL-cholesterol availability and cholesteryl ester turnover. Cancer Metab 2022;10(1):1 doi 10.1186/s40170-021-00278-1.

54. Liang P, Henning SM, Grogan T, Elashoff D, Said J, Cohen P, et al. Effect of omega-3 fatty acid diet on prostate cancer progression and cholesterol efflux in tumor-associated macrophages-dependence on GPR120. Prostate Cancer Prostatic Dis 2023 doi 10.1038/s41391-023-00745-4.

55. Sharma B, Agnihotri N. Role of cholesterol homeostasis and its efflux pathways in cancer progression. J Steroid Biochem Mol Biol 2019;191:105377 doi 10.1016/j.jsbmb.2019.105377.

56. Krycer JR, Phan L, Brown AJ. A key regulator of cholesterol homoeostasis, SREBP-2, can be targeted in prostate cancer cells with natural products. Biochem J 2012;446(2):191–201 doi 10.1042/BJ20120545.

57. Swinnen JV, Ulrix W, Heyns W, Verhoeven G. Coordinate regulation of lipogenic gene expression by androgens: evidence for a cascade mechanism involving sterol regulatory element binding proteins. Proc Natl Acad Sci U S A 1997;94(24):12975–80 doi 10.1073/pnas.94.24.12975.

58. Wang X, Sun B, Wei L, Jian X, Shan K, He Q, et al. Cholesterol and saturated fatty acids synergistically promote the malignant progression of prostate cancer. Neoplasia 2022;24(2):86–97 doi 10.1016/j.neo.2021.11.004.

59. Ding X, Zhang W, Li S, Yang H. The role of cholesterol metabolism in cancer. Am J Cancer Res 2019;9(2):219–27.

60. Antonarakis ES, Lu C, Wang H, Luber B, Nakazawa M, Roeser JC, et al. AR-V7 and resistance to enzalutamide and abiraterone in prostate cancer. N Engl J Med 2014;371(11):1028–38 doi 10.1056/NEJMoa1315815.

61. Watson PA, Arora VK, Sawyers CL. Emerging mechanisms of resistance to androgen receptor inhibitors in prostate cancer. Nat Rev Cancer 2015;15(12):701–11 doi 10.1038/nrc4016.

62. Abida W, Cyrta J, Heller G, Prandi D, Armenia J, Coleman I, et al. Genomic correlates of clinical outcome in advanced prostate cancer. Proc Natl Acad Sci U S A 2019;116(23):11428–36 doi 10.1073/pnas.1902651116.

63. Lee JH, Hong J, Zhang Z, de la Pena Avalos B, Proietti CJ, Deamicis AR, et al. Regulation of telomere homeostasis and genomic stability in cancer by N (6)-adenosine methylation (m(6)A). Sci Adv 2021;7(31) doi 10.1126/sciadv.abg7073.

